# Enhancing Medical Image Segmentation through Negative Sample Integration: A Study on Kvasir-SEG and Augmented Datasets

**DOI:** 10.1101/2025.10.14.682445

**Authors:** Imon Mahbub, Ahmed Zoha Zim, Talha Bin Imran, K. M. Fatin Mesbah, Md Jobayer, Md. Mehedi Hasan Shawon

**Affiliations:** BSRM School of Engineering, BRAC University, Dhaka, Bangladesh; Department of Biomedical Engineering, Linköping University, Sweden

## Abstract

Colorectal cancer (CRC) remains a leading cause of cancer-related mortality worldwide, with early and accurate detection being critical for improving patient outcomes. Automated image segmentation using deep learning has emerged as a transformative tool for identifying colorectal abnormalities in medical imaging. This study conducts a comparative analysis of three prominent deep learning architectures—U-Net, SegNet, and ResNet—for colorectal cancer image segmentation, evaluating their performance on a custom dataset comprising 1,800 images (1,000 polyp images from the Kvasir-SEG dataset and 800 polyp-free images from the WCE Curated Colon Dataset). The dataset was preprocessed to a uniform resolution of 256 × 256 pixels and partitioned into training, validation, and test sets. Quantitative and qualitative results demonstrate that U-Net outperforms SegNet and ResNet, achieving superior segmentation accuracy (validation accuracy of 0.95) and robustness, particularly when trained on datasets that include negative samples. SegNet showed the sign of overfitting and delivered unstable results, while ResNet struggled to generalize effectively. The integration of negative images improved specificity by decreasing false positive rates. Overall, the results demonstrate U-net as the most efficient in precise polyp segmentation, providing significant implications for robust diagnostic system development.

## Introduction

CRC represents a significant global health concern, listed among the foremost causes of cancer incidence and death. GLOBOCAN 2018 data shows that the CRC disease accounts for an estimated 1.8 million new cases and 850,000 deaths each year worldwide [1]. Although older adults over 65 carry the greatest risk, younger populations are increasingly affected due to genetic predisposition, lifestyle habits, and rising obesity rates [2]. Early and accurate detection of colorectal abnormalities is therefore essential to improving survival rates. In clinical settings, endoscopic imaging plays a pivotal role in detecting polyps and tumors, yet manual interpretation is subjective, time-consuming, and highly dependent on clinician expertise [3].

Deep learning (DL) has emerged as a transformative solution for automated medical image segmentation. Convolutional neural networks (CNNs), and architectures such as U-Net [10], SegNet [6], and ResNet-based variants [15], have demonstrated outstanding performance in extracting spatial and structural features for delineating pathological regions [4, 25]. Despite this progress, most segmentation studies emphasize only positive samples (images containing polyps), overlooking negative cases that are equally common in real-world practice. This bias can lead to models that perform well on benchmark datasets but generate high false-positive rates in clinical deployment. Integrating polyp-free samples into training has the potential to improve model robustness and clinical applicability, yet this remains underexplored [39, 40].

Moreover, patient-related factors such as obesity contribute to nearly 10% of CRC cases and influence tumor morphology and tissue microenvironment through inflammatory and hormonal mechanisms [7, 9]. Automated segmentation tools that can adapt to such variations are critical for reliable diagnostic support. However, existing comparative studies seldom evaluate model performance under these clinically realistic conditions.

To address this gap, this study systematically benchmarks three widely adopted DL architectures—U-Net, SegNet, and ResNet—using a uniquely composed dataset. The dataset combines polyp images with expert-annotated masks from Kvasir-SEG [27] and polyp-free images from the WCE Curated Colon Disease Dataset [26]. By explicitly integrating negative samples, we aim to investigate their effect on segmentation performance and establish a more realistic benchmark for colorectal cancer image analysis. The findings of this work are expected to provide deeper insights into building robust, clinically relevant DL-based segmentation models that enhance early CRC detection.

### Related Work

Among the various deep learning (DL) architectures, models such as U-Net [10], SegNet [6], and ResNet-based networks [8] have consistently demonstrated strong performance in medical image analysis due to their robust feature extraction capabilities and specialized segmentation designs. The U-Net model, characterized by its encoder–decoder symmetry and skip connections, is particularly effective in maintaining spatial detail thereby supporting accurate localization of boundaries in medical images [12]. In contrast Segnet, another encoder–decoder approach, achieves efficiency by reusing max-pooling indices during the upsampling stage, which makes it favorable when computational resources are constrained [15]. On the other hand, ResNet-based models address the vanishing gradient challenge, facilitating deeper architectures that can extract sophisticated features [3, 15].

In addition to the base models, numerous extensions and adaptations have been introduced to meet the unique demands of specific tasks. Edge U-Net [35] integrates edge detection into the U-Net framework to enhance boundary precision, which is particularly valuable in colorectal image segmentation where delineating polyp edges is critical. U-Net++ [5], a nested variant of U-Net, improves multi-scale feature extraction and boundary delineation, though its increased computational complexity limits real-time applications. ResUNet++ [17] further enhances the U-Net structure with residual units, squeeze-and-excitation blocks, and atrous spatial pyramid pooling, demonstrating improved segmentation of complex structures such as colorectal polyps. Similarly, Focus U-Net [16], which incorporates dual attention gates, shows strong performance on colonoscopy datasets but was evaluated without negative samples, raising questions about its robustness in real-world scenarios.

Lightweight architectures have also been explored to balance segmentation accuracy and real-time applicability. HarDNet-MSEG [18], employing HarDNet as its backbone, achieves mean Dice scores above 0.9 with inference speeds exceeding 86 FPS. While highly efficient, its evaluation, like many other approaches, was limited to polyp-containing datasets, leaving its performance on negative samples untested. This omission is critical since non-polyp frames dominate real-world colonoscopy procedures, and failure to address them increases the likelihood of false positives.

Beyond encoder–decoder variants, Fully Convolutional Networks (FCNs) and other CNN-based methods have been comprehensively reviewed in the literature for their contributions to medical image segmentation [3, 4, 21]. Domain-specific adaptations, such as CNNs designed for skin lesion segmentation [40], highlight how specialized architectures can improve detection accuracy and provide insights into potential modifications for colorectal cancer segmentation. Multi-scale CNNs [31], originally proposed for lung nodule detection, also demonstrate the importance of capturing features at multiple resolutions—an approach directly transferable to polyp segmentation, where lesion sizes vary considerably.

Generative Adversarial Networks (GANs) have been explored for medical image synthesis, particularly in CT and MRI domains, to support data augmentation and anonymization [39]. While GANs can enrich datasets and potentially improve segmentation training [14], their limitations include high computational costs and variable image realism, which hinder clinical adoption. Hybrid approaches such as ReGANs [3], which integrate semi-automatic region growing with DL, illustrate the potential of semi-supervised learning in cases where partial manual guidance is available. Furthermore, optimization strategies such as incorporating attention mechanisms and lightweight designs into U-Net [13] demonstrate how architectural refinements can boost segmentation accuracy on complex datasets.

Beyond the medical domain, deep learning applications in plant disease detection [14] and hybrid models for heart disease prediction [3] provide additional insights. These studies emphasize the role of large-scale datasets, data augmentation, and multi-source data fusion—principles that could be adapted to colorectal imaging by integrating diverse data types (e.g., colonoscopy frames and clinical metadata) to improve diagnostic accuracy.

Overall, the reviewed literature highlights that while U-Net and its variants, along with lightweight and hybrid models, have significantly advanced medical image segmentation, a recurring limitation is the lack of evaluation on datasets containing negative samples. Addressing this gap is crucial for ensuring robust performance in real-world colorectal cancer screening, where distinguishing polyp-free frames is as important as accurately segmenting polyp-containing ones.

## Methodology

Following dataset preparation, three deep learning architectures—U-Net, SegNet, and a ResNet-based model were employed for colorectal image segmentation. As illustrated in Fig 1 and Fig 2, the methodology began with the construction of a custom dataset that combined polyp images with corresponding ground truth masks.

**Fig 1.**
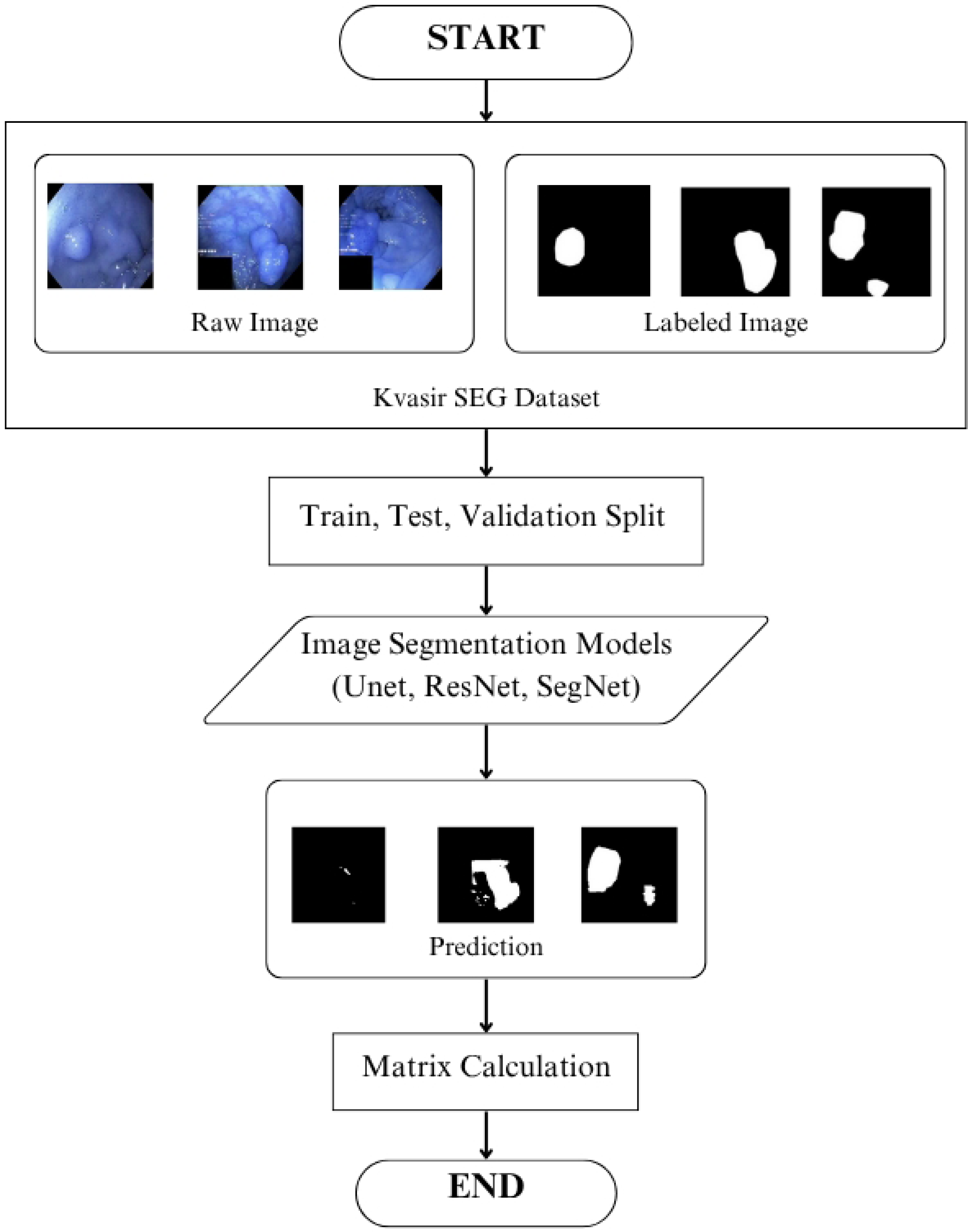
Overview of the proposed methodology. The Kvasir-SEG dataset is preprocessed and divided into training, validation, and testing sets. Segmentation models (U-Net, ResNet, SegNet) are trained and evaluated, followed by prediction generation and quantitative performance assessment.

**Fig 2.**
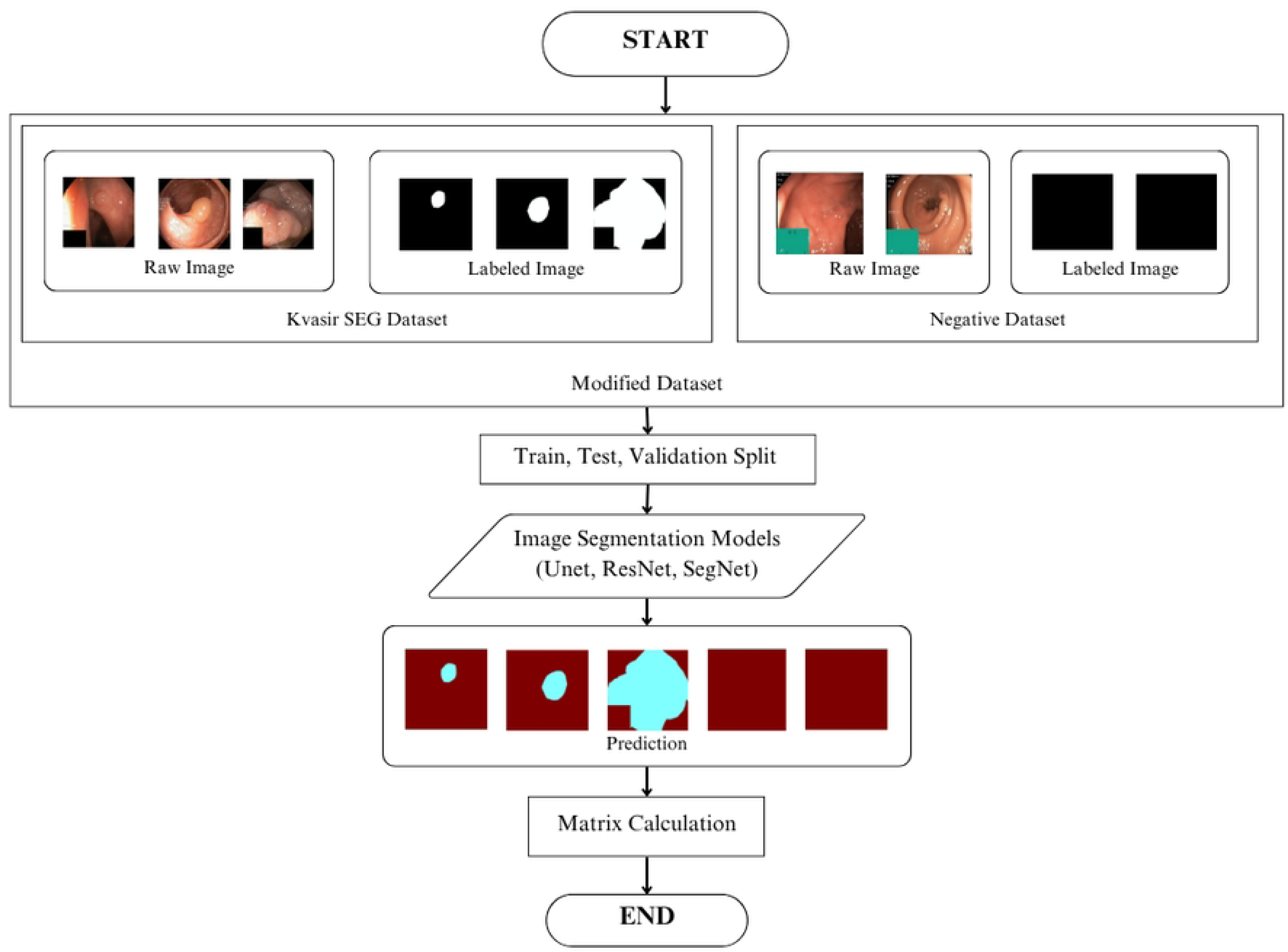
Illustration of the modified dataset pipeline. In addition to the original Kvasir-SEG dataset, negative samples (non-polyp images) are incorporated to create a more robust dataset. The combined dataset is split into training, validation, and testing sets. Segmentation models (U-Net, ResNet, SegNet) are trained on this modified dataset, followed by prediction generation and performance evaluation through quantitative metric calculations.

Fig 1 represents the baseline methodology, which utilizes only the original Kvasir-SEG dataset [27]. In this approach, raw colonoscopy images and their corresponding labeled masks are preprocessed and split into training, validation, and test subsets. Segmentation models (U-Net, ResNet, SegNet) are then trained, predictions are generated, and performance is quantitatively assessed.

Fig 2 illustrates the modified dataset pipeline, where both positive (polyp) and negative (polyp-free) [26] samples are integrated to improve model robustness and specificity. The combined dataset undergoes the same split into training, validation, and testing subsets, followed by training with the three architectures. Predictions are generated for the unseen test set, and quantitative metrics (e.g., Dice coefficient, IoU, precision, recall) are calculated to evaluate segmentation performance.

This systematic design allowed the models to not only learn accurate polyp segmentation but also to differentiate polyp-free cases, thereby enhancing specificity and real-world applicability. The comparative evaluation of the three architectures provided empirical insights into their effectiveness for colorectal abnormality detection and segmentation.

### Dataset

The study employed a carefully constructed dataset that combined two primary sources to ensure a comprehensive evaluation. The Kvasir-SEG dataset [27] comprises 1,000 high-resolution endoscopic images (ranging from 720×576 to 1920×1080 pixels), each accompanied by manually annotated polyp masks that expert gastroenterologists have rigorously validated. This collection represented diverse polyp morphologies, including flat, sessile, and pedunculated types, captured under varying illumination conditions to enhance real-world applicability. Complementing this, the Curated Colon Dataset [26] provided 800 normal colonoscopy images (averaging 500 × 500 pixels) depicting healthy mucosal patterns. These negative samples were carefully selected to represent various anatomical landmarks, such as the ileocecal valve and hepatic flexure, while maintaining a balanced distribution between clean mucosal views and those with residual fluid or air bubbles, which are common artifacts in clinical practice. A robust preprocessing pipeline standardized all images to 256×256 pixels using bicubic interpolation, followed by histogram equalization to normalize illumination variations. To further enhance model generalization, the study implemented comprehensive data augmentation, including random rotations (±15°), horizontal/vertical flips (50% probability), color jitter (±10% brightness/contrast adjustments), and controlled Gaussian noise injection (*σ* = 0.01) to simulate real-world imaging variations while preserving diagnostic features.

### Model Architecture and Specification

Three deep learning architectures: U-Net, SegNet and ResNet were trained and evaluated under identical conditions for fair comparison. Each models code is available on GitHub https://github.com/imonM007/MedKvSet).

### U-Net

The U-Net model, introduced by Ronneberger et al. [10], stands as a cornerstone in the field of biomedical image segmentation. Its importance stems from its unique symmetric encoder–decoder architecture, which is specifically designed to excel with the challenges posed by medical data. The encoder, a contracting path composed of repeated convolutional blocks (each with two 3×3 convolutional layers followed by a ReLU activation) and 2×2 max-pooling layers for downsampling, progressively captures high-level semantic features, such as the general presence of a polyp. The decoder, or expansive path, then upsamples these features using transposed convolutions (also known as deconvolutional layers) to generate a segmentation map of the same size as the original image. The most critical innovation of U-Net is its use of “skip connections.” These concatenation-based skip connections directly link the high-resolution feature maps from each encoder stage to the corresponding decoder stage at the same spatial resolution. This fusion of features allows the model to leverage both global contextual information from the deeper layers and fine-grained spatial details from the earlier layers. This is particularly vital in medical imaging, where the precise boundaries and intricate textures of abnormalities, such as polyps, must be accurately delineated for a correct diagnosis. The ability to retain this localization information allows U-Net to produce highly accurate and detailed segmentation masks, making it a powerful tool even when training on limited datasets, making a common practice to use the model in various field such as crack detection, satellite imagery and autonomous driving [11], [12], [13].

### SegNet

A SegNet-based convolutional neural network was implemented in this study for medical image segmentation. SegNet has been established widely as an effective semantic segmentation model, particularly suited for pixel-wise classification tasks that demand accurate boundary representation. Although it was initially designed for scene understanding, SegNet has been adapted for numerous dense prediction tasks, including applications in medical imaging [6]. SegNet architecture follows an encoder–decoder design with the encoder consisting of 13 convolutional layers derived from VGG16, arranged into modules of convolution, batch normalization, ReLU activation, and max-pooling. Pooling indices are retained during the downsampling stage and it was reused by the decoder for efficient upsampling without additional parameters. The decoder mirrors this structure, employing the stored indices to upsample feature maps which is followed by convolutions to refine outputs and it concludes with a softmax classifier to generate pixel-wise probabilities. With about 29.5 million trainable parameters, SegNet maintains memory efficiency, supports arbitrarily sized RGB inputs, and produces detailed segmentation outputs making it particularly advantageous in resource-limited environments [6], [19], [20], [21].

### ResNet Model

The disease recognition system in this study leverages the Residual Network (ResNet) architecture, specifically ResNet18, to process paired images (original images and their corresponding masks) for binary classification, drawing from foundational advancements in deep learning. ResNet, introduced by He et al. [15], is celebrated for its use of residual connections, which mitigate the vanishing gradient problem by adding a layer’s input directly to its output, enabling effective training of very deep models. These residual blocks facilitate the extraction of increasingly abstract and complex features, making ResNet a powerful feature extractor for medical image analysis. The use of pretrained weights from large-scale datasets such as ImageNet, as described by Deng et al. [22] and Russakovsky et al. [23], supports transfer learning by adapting general visual representations to specialized medical domains, improving performance on limited datasets. To enhance generalization, data augmentation techniques like random flips, rotations, and color jitter are applied, inspired by Szegedy et al. [24]. However, despite its strengths, a standard ResNet architecture faces challenges when extended to pixel-wise segmentation tasks. While residual connections reduce information loss during training, downsampling operations in deeper networks can compromise spatial detail, which is critical for precise segmentation masks. This tendency, coupled with the risk of overfitting on smaller medical datasets, indicates that architectural modifications or careful regularization are necessary for optimal performance in segmentation [25].

## Results

### U-Net Model Output Analysis

#### Training Behavior

Figure 6 shows the training and validation curves of U-Net on both datasets.

**Fig 3.**
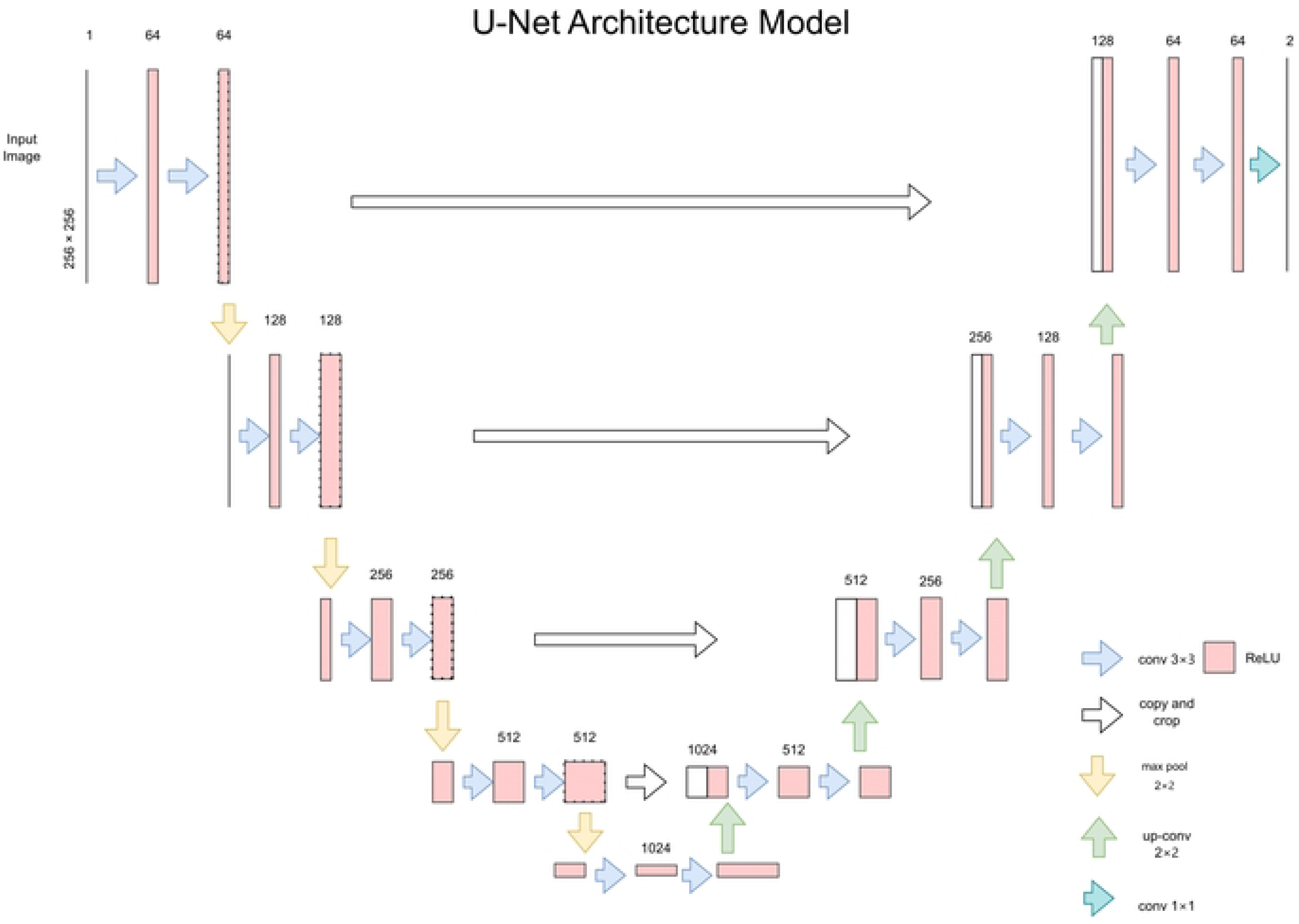
The figure represents the U-Net architecture, designed with a contracting path for feature representation and an expansive path for precise localization. The diagram highlights the combined role of convolution, pooling and up-convolution layer in providing precise segmentation outputs.

**Fig 4.**
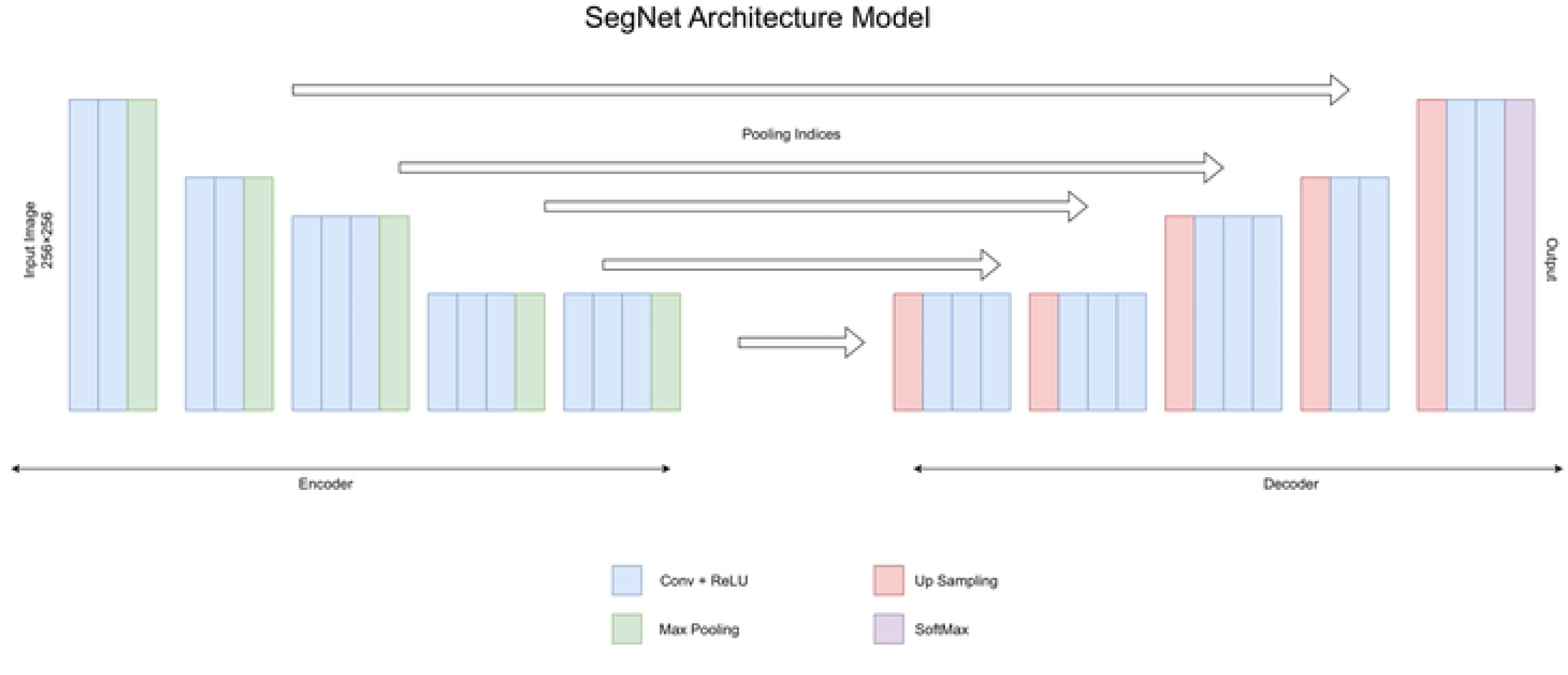
This figure represents the SegNet architecture, consisting of an encoder for feature extraction and a decoder for reconstructing the segmentation maps. The model utilizes convolution with ReLU, max pooling with stored indices, and up-sampling to map input images to their segmented outputs accurately.

**Fig 5.**
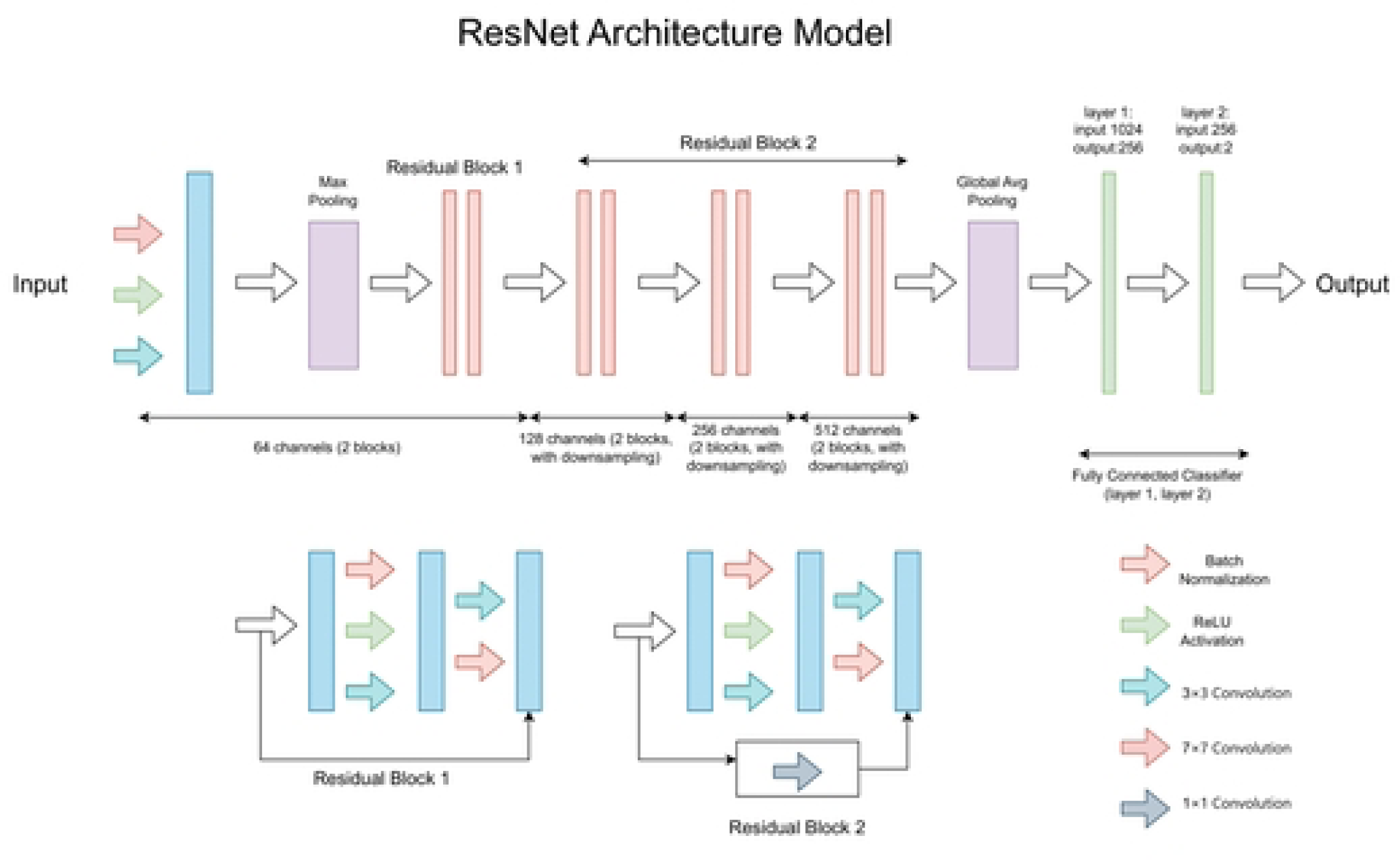
This diagram outlines the ResNet model’s structure, starting with an input layer and including max pooling and residual blocks with the increasing of channel depths. It features global average pooling and a fully connected classifier, enhanced by convolutions and normalization for effective image classification.

**Fig 6.**
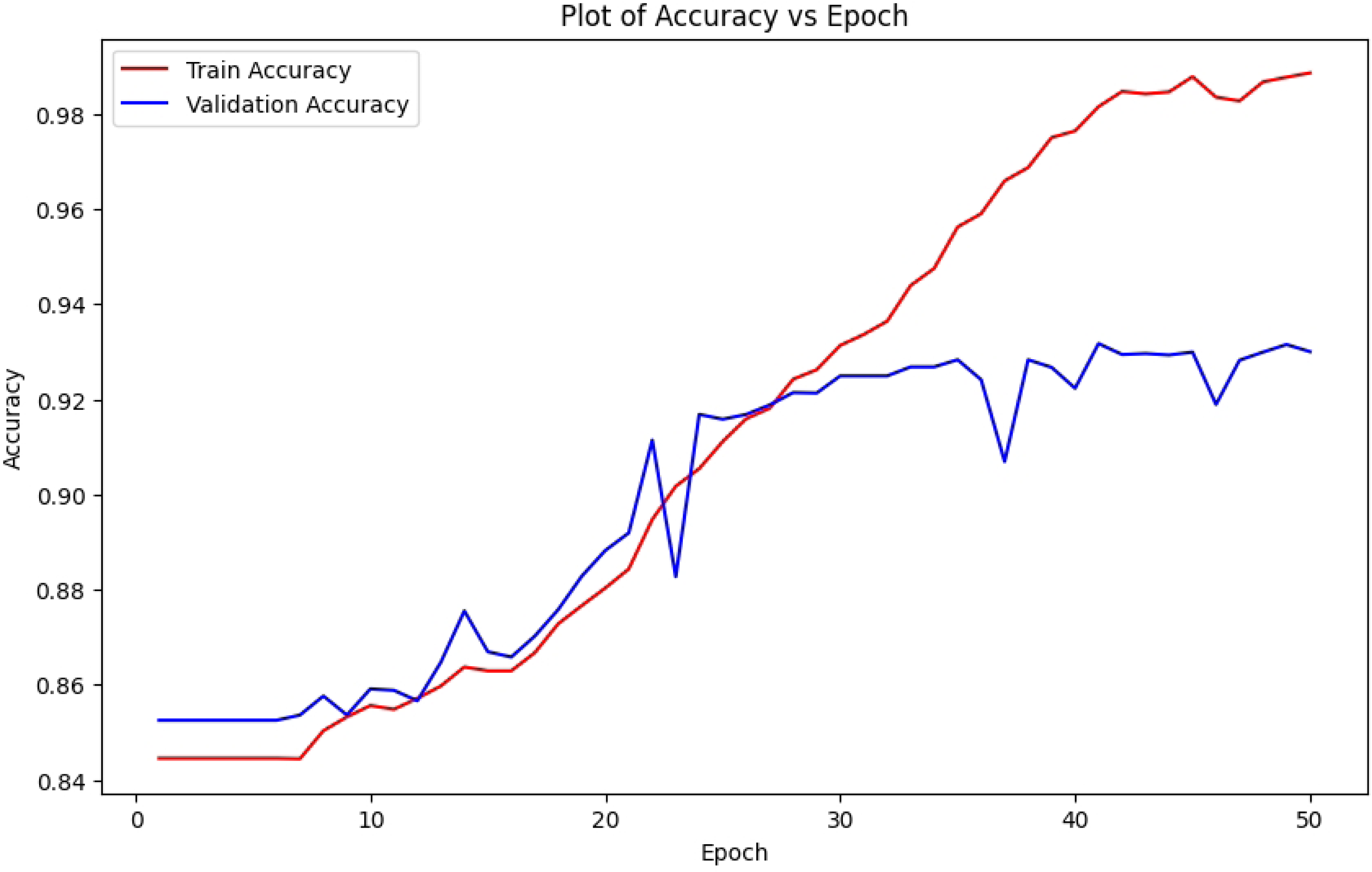
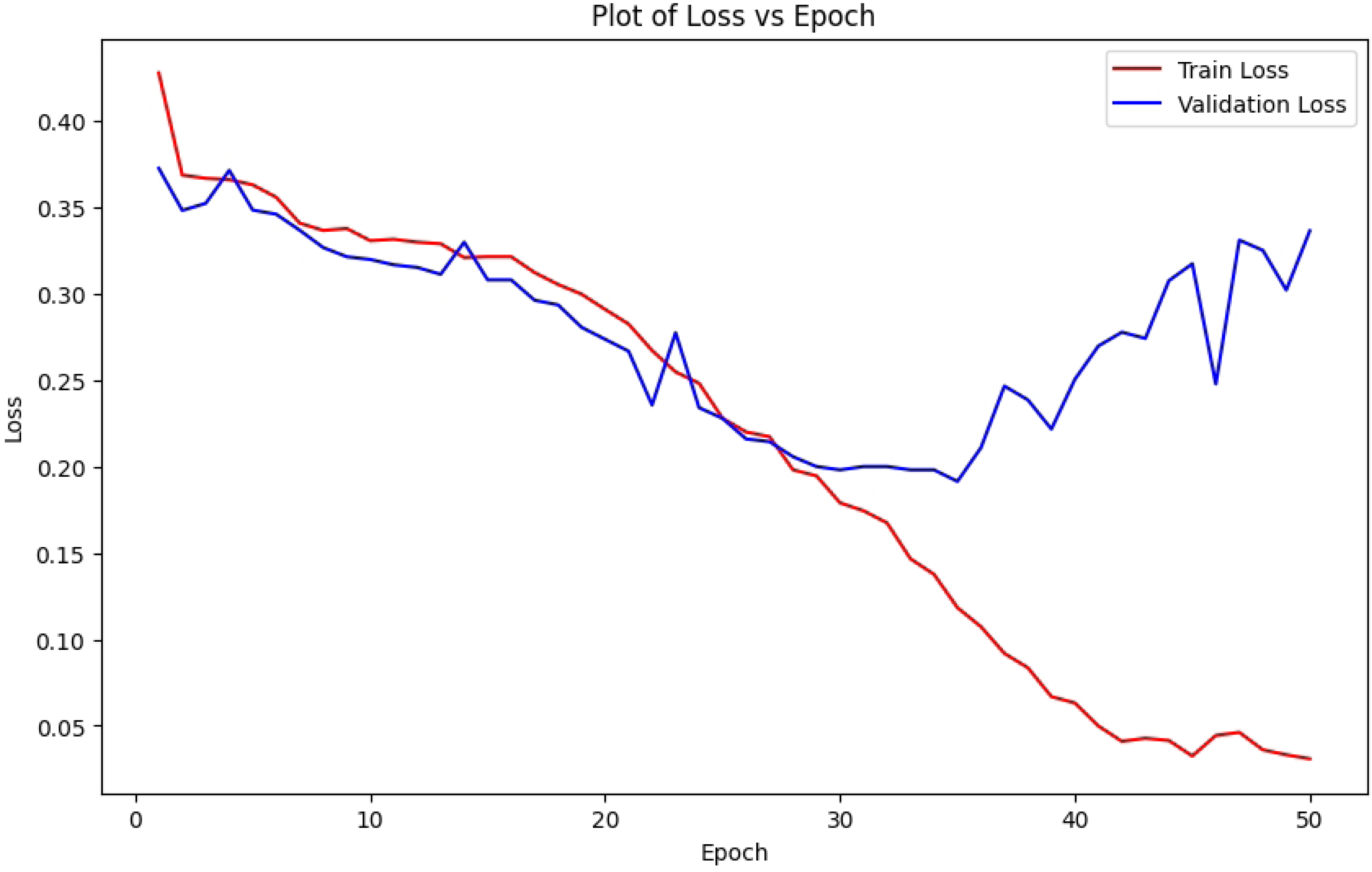
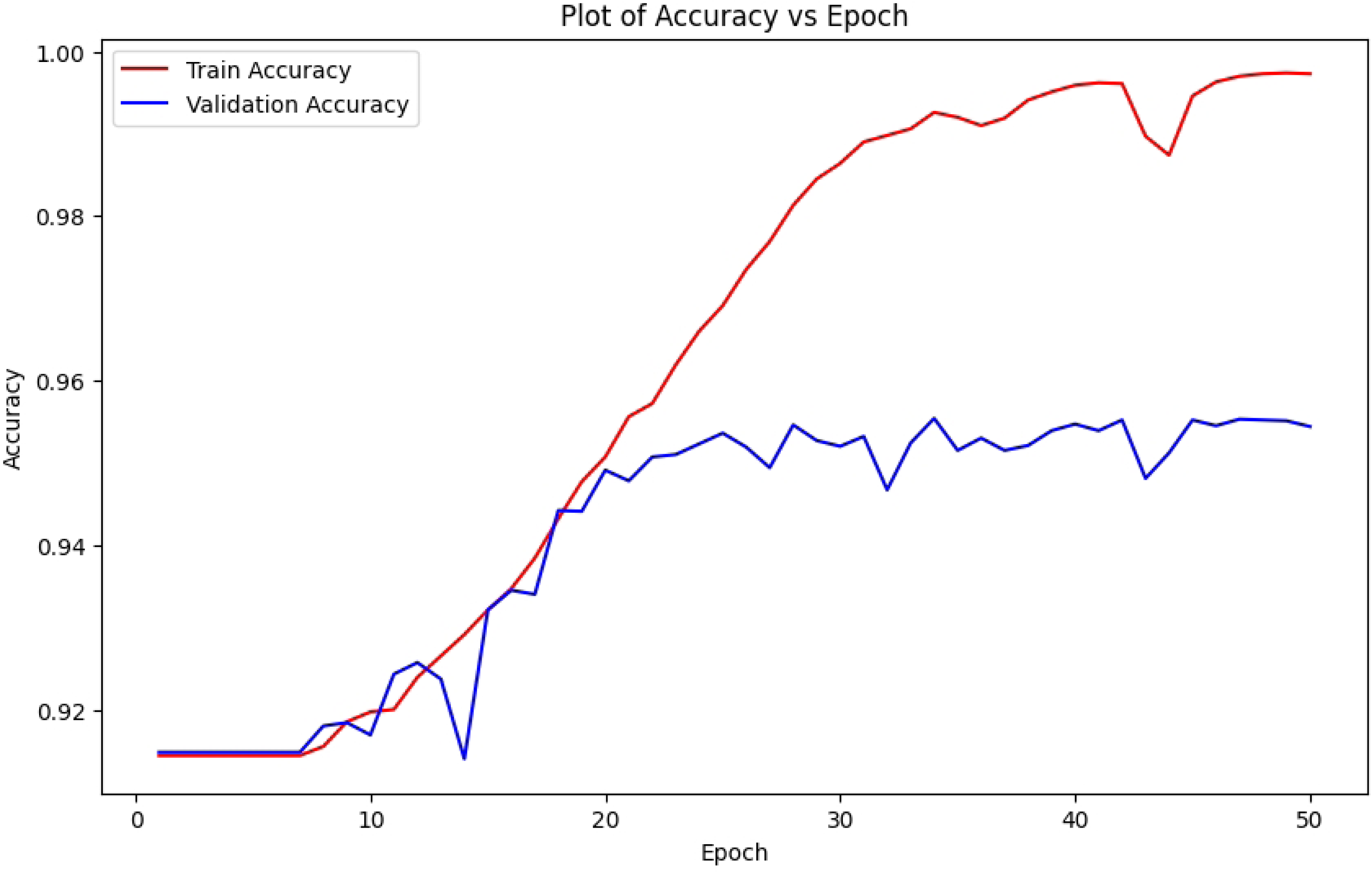
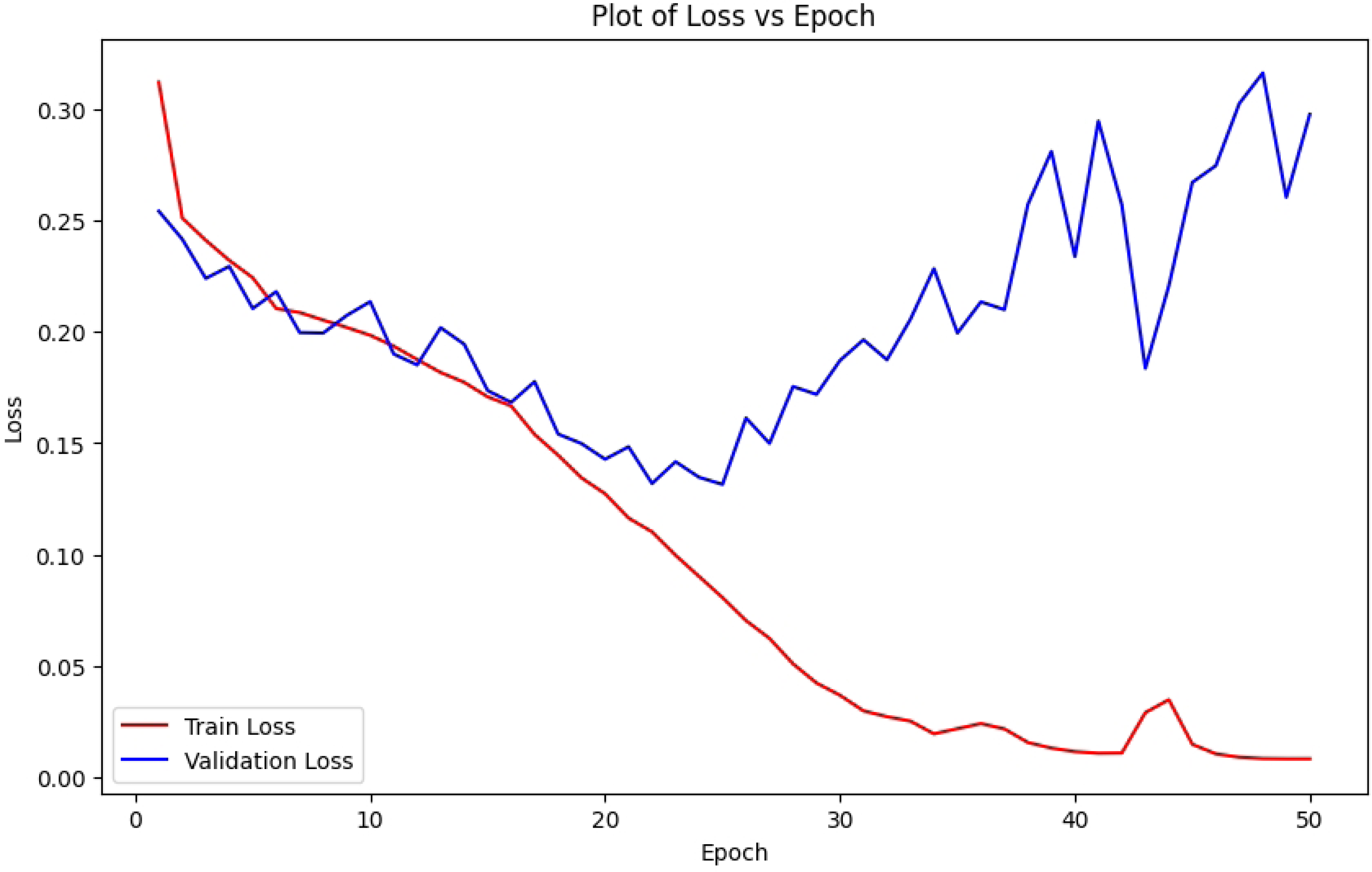
This set of graphs illustrates the direction of training and validation accuracy and loss over 50 epochs for the U-Net model, showing improved model performance and stability with datasets of 1000 and 1800 images.

The first two graphs—train and validation accuracy vs. epoch and train and validation loss vs. epoch in Figure 6 (a) (b)—illustrate U-Net training progress over 50 epochs. In the accuracy plot, training steadily rises from 0.84 to nearly 0.98, while validation improves to about 0.92 but fluctuates and declines slightly after epoch 30, suggesting overfitting. The loss plot shows training loss dropping from 0.40 to below 0.05, while validation loss decreases initially but rises sharply after epoch 30 to around 0.35, again indicating overfitting. These trends imply that while the model learns well, its generalization weakens after 30 epochs, warranting early stopping or regularization.

The next two graphs—train/validation accuracy vs. epoch and loss vs. epoch in Figure 6 (c) (d)—use the larger dataset. Training accuracy approaches 1.0, while validation accuracy plateaus at 0.94–0.96 after epoch 20 with fluctuations, suggesting overfitting. Training loss decreases from 0.30 to below 0.05, but validation loss, after an initial drop, rises to 0.25 beyond epoch 20, confirming weaker generalization. Overall, strong training performance contrasts with divergence in validation metrics after 20 epochs, indicating that early stopping or regularization could improve generalization.set.

(a) Train and Validation Accuracy VS Epoch (1000 Image)
(b) Train and Validation Loss VS Epoch (1000 Image)
(c) Train and Validation Accuracy VS Epoch (1800 Image)
(d) Train and Validation Loss VS Epoch (1800 Image)

### Qualitative Output

Figure 7 presents qualitative outputs from U-Net before and after adding negative data. Each sample includes:

**Fig 7.**
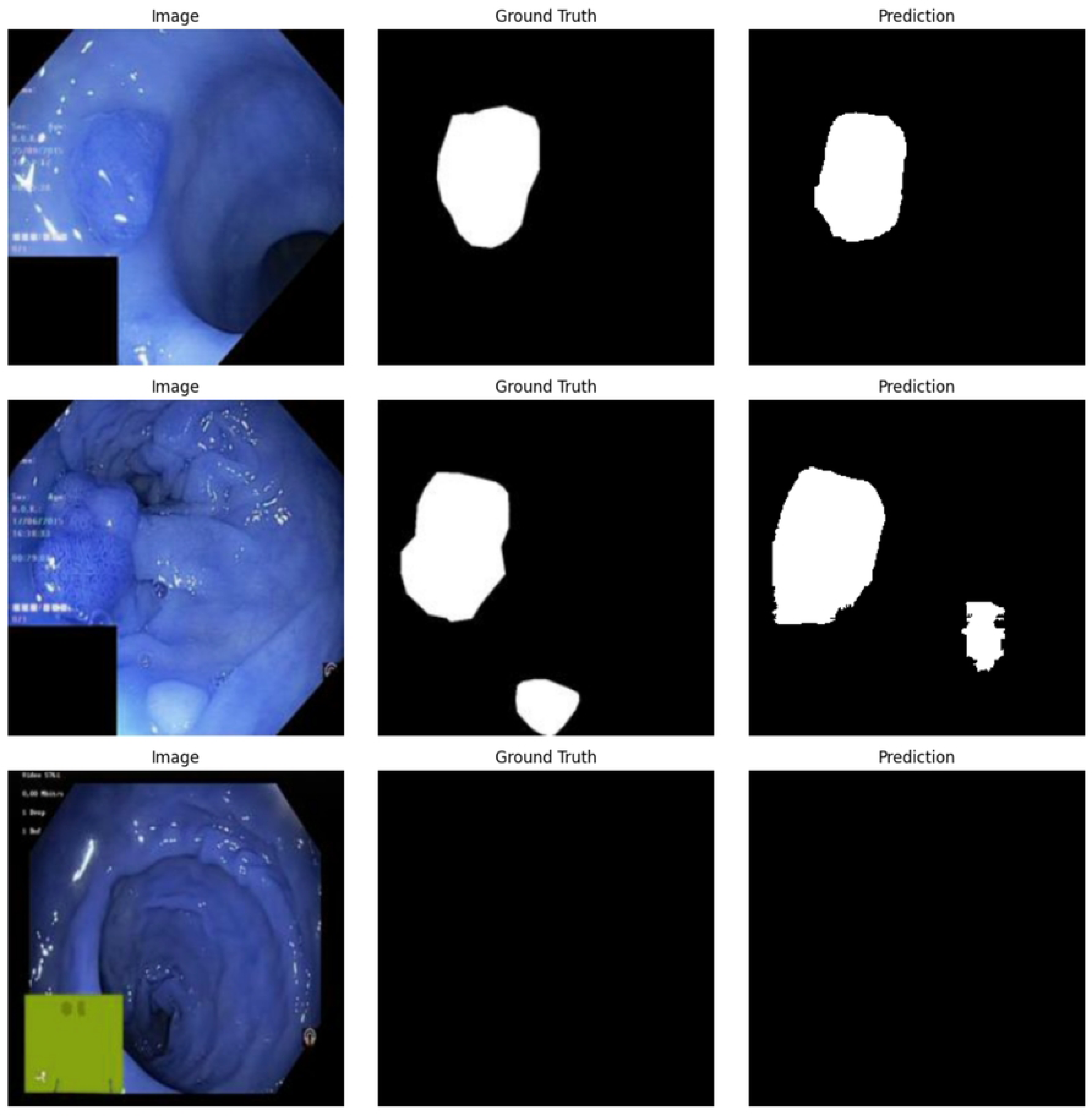
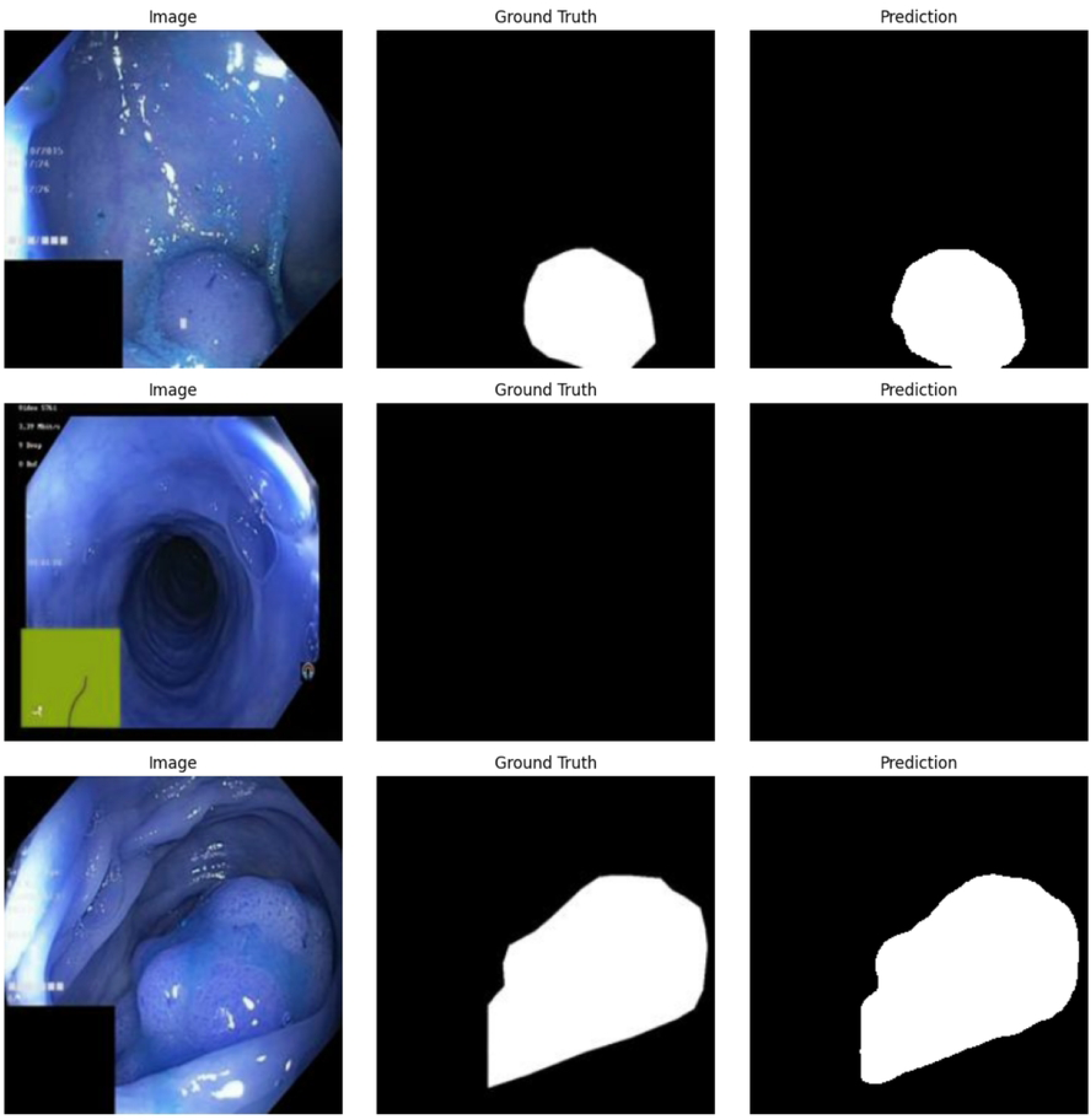
This image collection for the U-Net model first showcases the effect images, followed by their ground truth and predicted segmentations, highlighting moderate to improved accuracy and alignment with actual regions as the dataset expands from 1000 to 1800 images.

The predicted masks closely resemble the ground truth masks, effectively capturing polyp shapes and locations. For the first two images in Figure 7 (a), the predictions align well with the single polyp regions, though minor boundary irregularities are visible. The third image shows a partial prediction, missing a secondary polyp, indicating some limitation in detecting multiple or smaller wounds. Here in Figure 7 (b), the U-Net model, trained on a dataset of 1,800 images, including 800 negative cases, shows compelling results in medical image segmentation. A significant strength of the U-Net model lies in its ability to correctly classify negative images, accurately predicting no segmentation when no target is present, which is important for minimizing false positives in the training data. In the output prediction (b) of Figure 7, for the first and last images, the model perfectly identified the polyp and gives a predicted area based on the ground truth. Also, in the second image, there is no polyp and in the prediction, there is no mask of polyp, so the model performed accurately in predicting the polyp images.

The U-Net model has shown stable training behavior, achieving high segmentation accuracy in the dataset due to its effective encoder-decoder architecture. Also, the U-Net model achieves a high dice coefficient, but its performance drops in complex polyps with irregular shapes and varying textures, which indicates the limitations in capturing various morphological features [35].

(a) Prediction after training with 1000 Image
(b) Prediction after training with 1800 Image

### SegNet Model Output Analysis

#### Training Behavior

Figure 8 shows the SegNet training and validation curves on both datasets.

**Fig 8.**
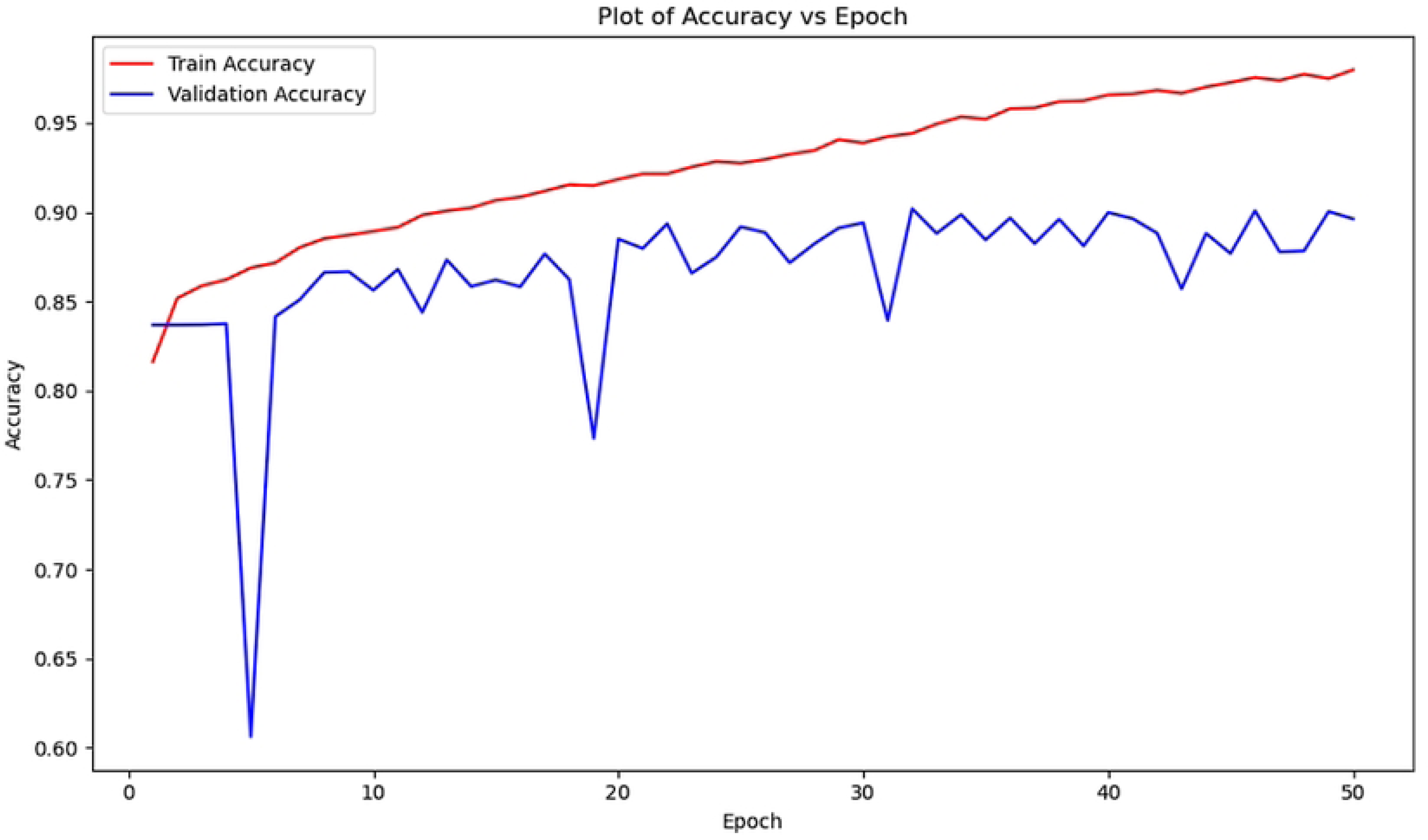
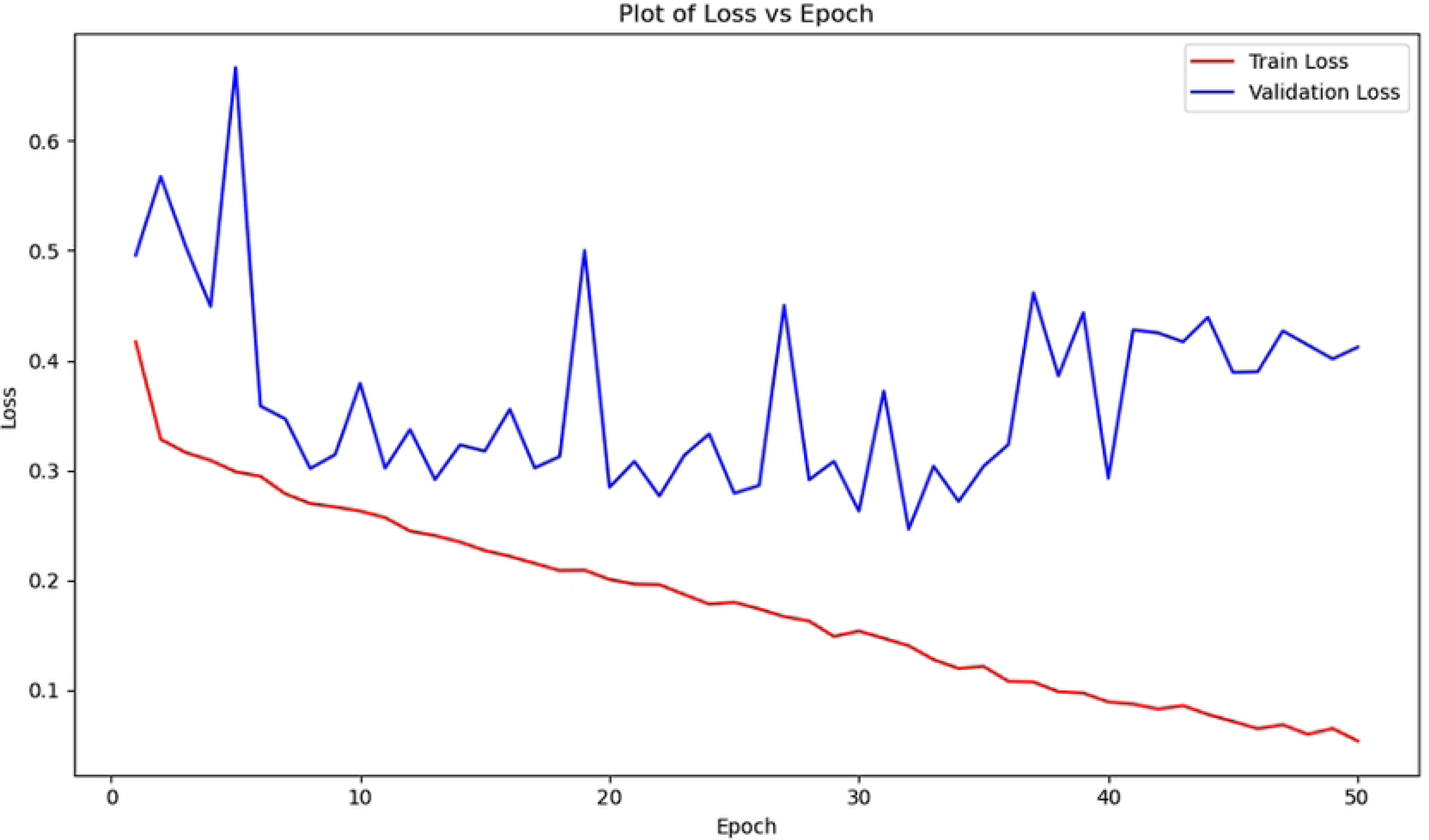
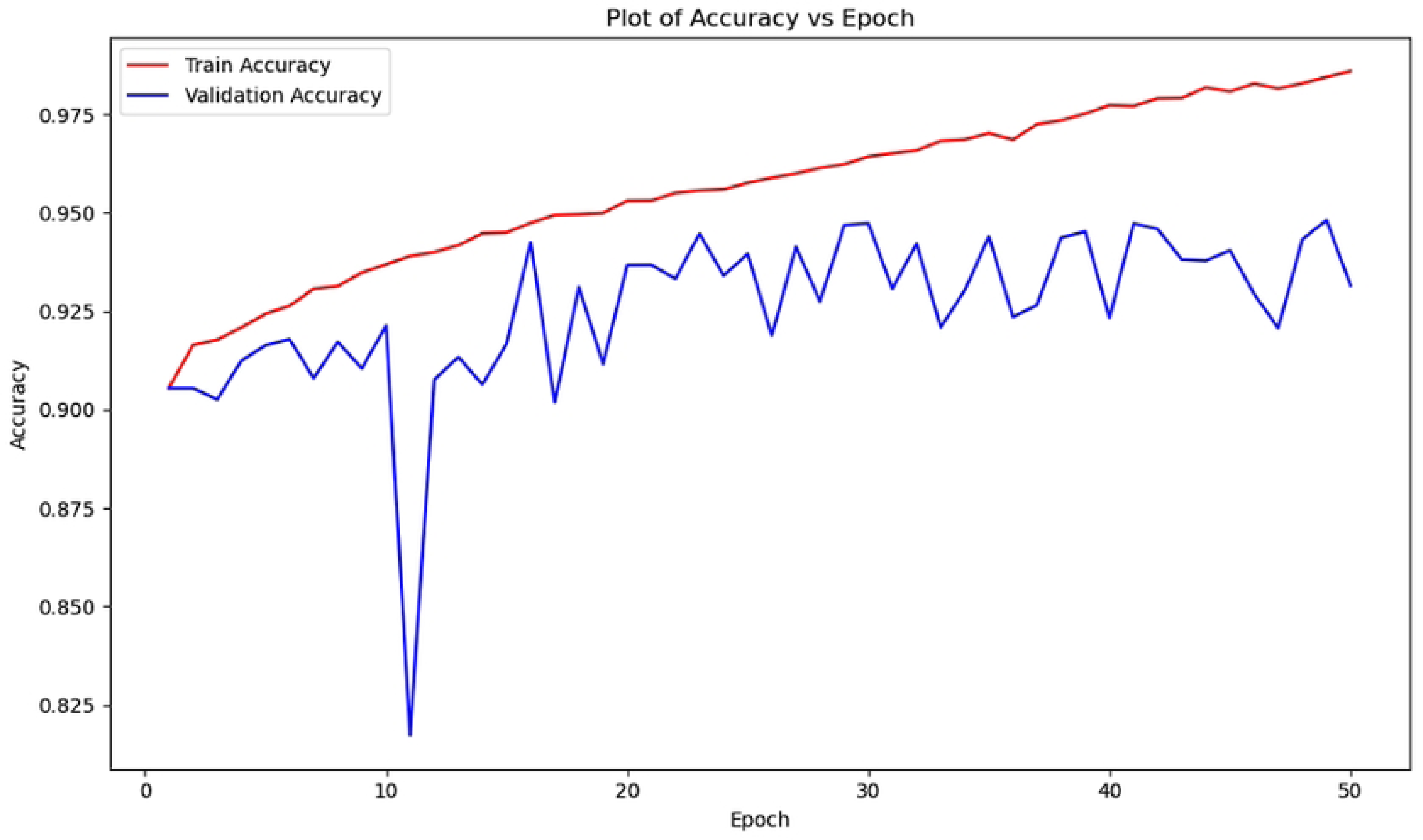
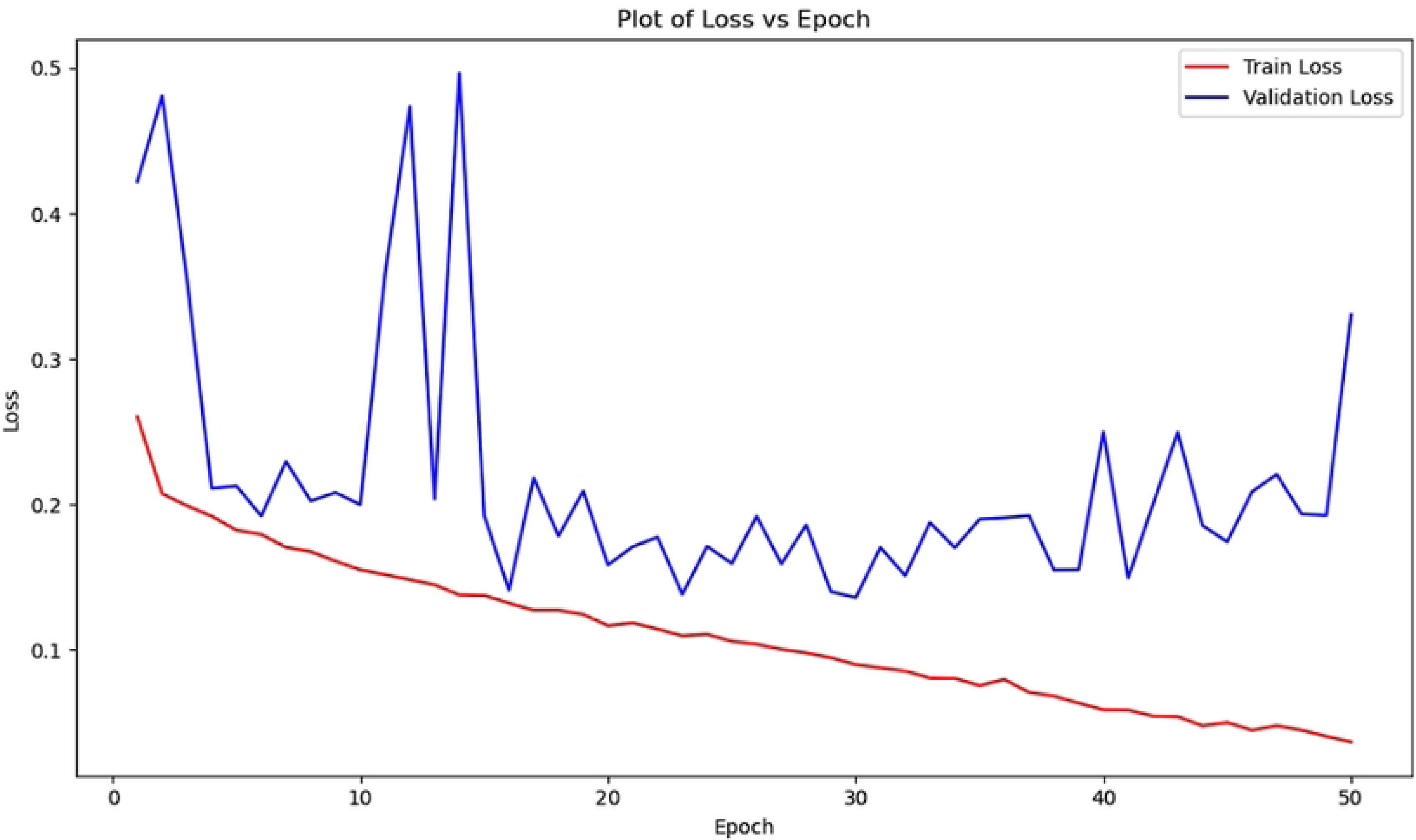
This set of graphs illustrates the trends in training and validation accuracy and loss over 50 epochs, showing the SegNet model’s learning progress and improved generalization with datasets of 1000 and 1800 images, where accuracy rises and loss decreases with reduced overfitting.

The first two images in 8 (a) (b) represent training/validation accuracy vs. epoch and loss vs. epoch with the 1000-image dataset. In Figure (a), training accuracy rises from 0.82 to 0.97 by epoch 50 with minor oscillations, while validation accuracy drops sharply from 0.84 to 0.60 at epoch 5, then fluctuates between 0.85–0.90 with peaks and troughs near epochs 5, 19, and 31. This instability, contrasted with steadily improving training accuracy, suggests overfitting due to dataset size or segmentation complexity. In Figure (b), training loss declines from 0.40 to 0.05, but validation loss spikes to 0.50 around epoch 5, later fluctuating between 0.30–0.50 with peaks near epochs 15 and 25, highlighting unstable performance despite gradual stabilization.

Figures 8 (c) (d) show results with the expanded dataset including negative samples.

In Figure (c), training accuracy climbs from 0.90 to 0.975 by epoch 50, while validation accuracy starts near 0.90, dips to 0.82 at epoch 11, then fluctuates between 0.925–0.950, reflecting reduced stability despite overall good performance. In Figure (d), training loss decreases smoothly from 0.25 to 0.05, but validation loss fluctuates heavily, peaking near 0.50 around epochs 3, 12, and 15 before stabilizing between 0.20–0.30. This divergence, with training loss minimized but validation loss higher, indicates overfitting and challenges in generalization introduced by negative samples.

(a) Train and Validation Accuracy VS Epoch (1000 Image)
(b) Train and Validation Loss VS Epoch (1000 Image)
(c) Train and Validation Accuracy VS Epoch (1800 Image)
(d) Train and Validation Loss VS Epoch (1800 Image)

### Qualitative Output

Figure 9 presents qualitative outputs from SegNet before and after adding negative data. Each sample includes:

**Fig 9.**
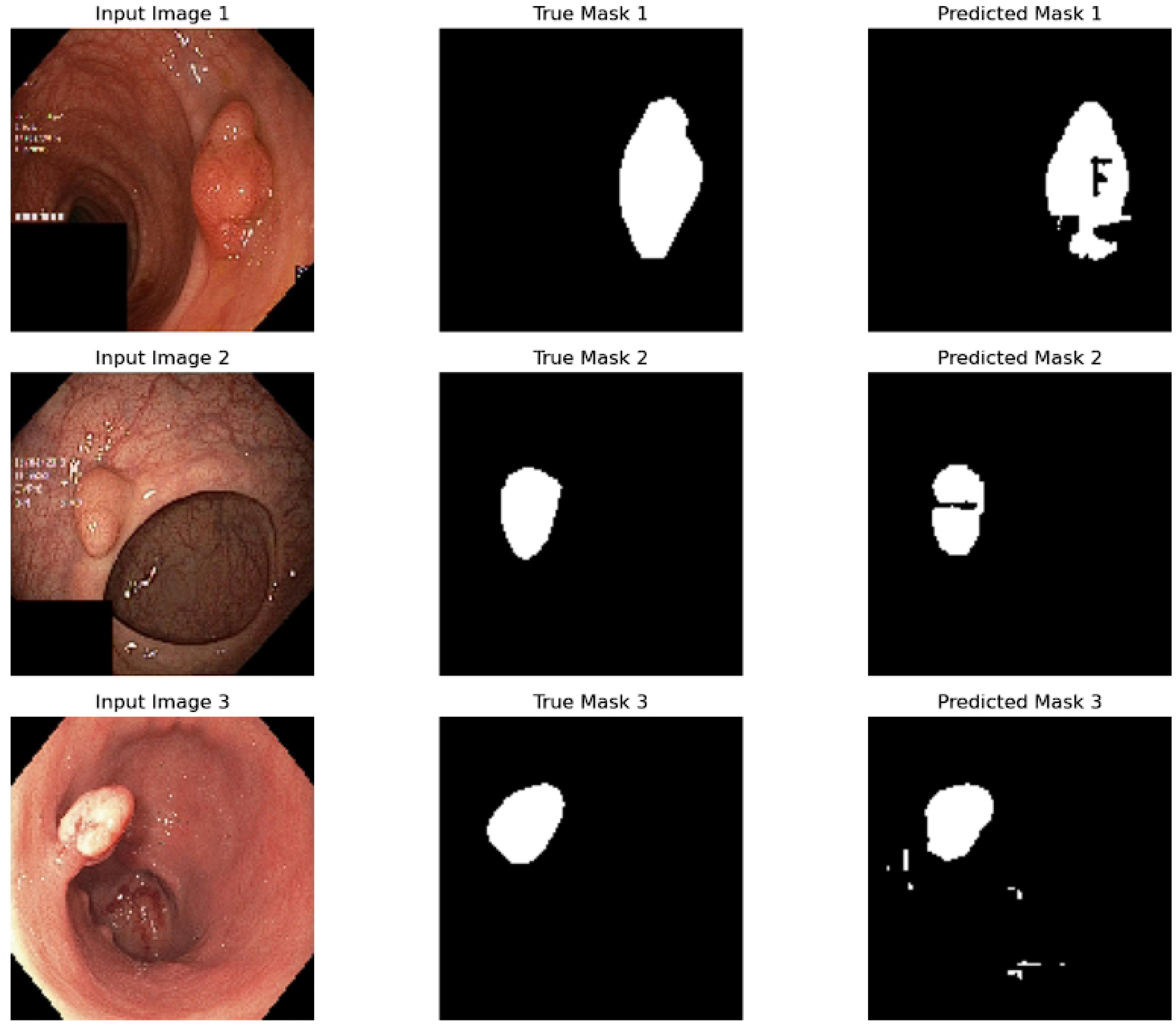
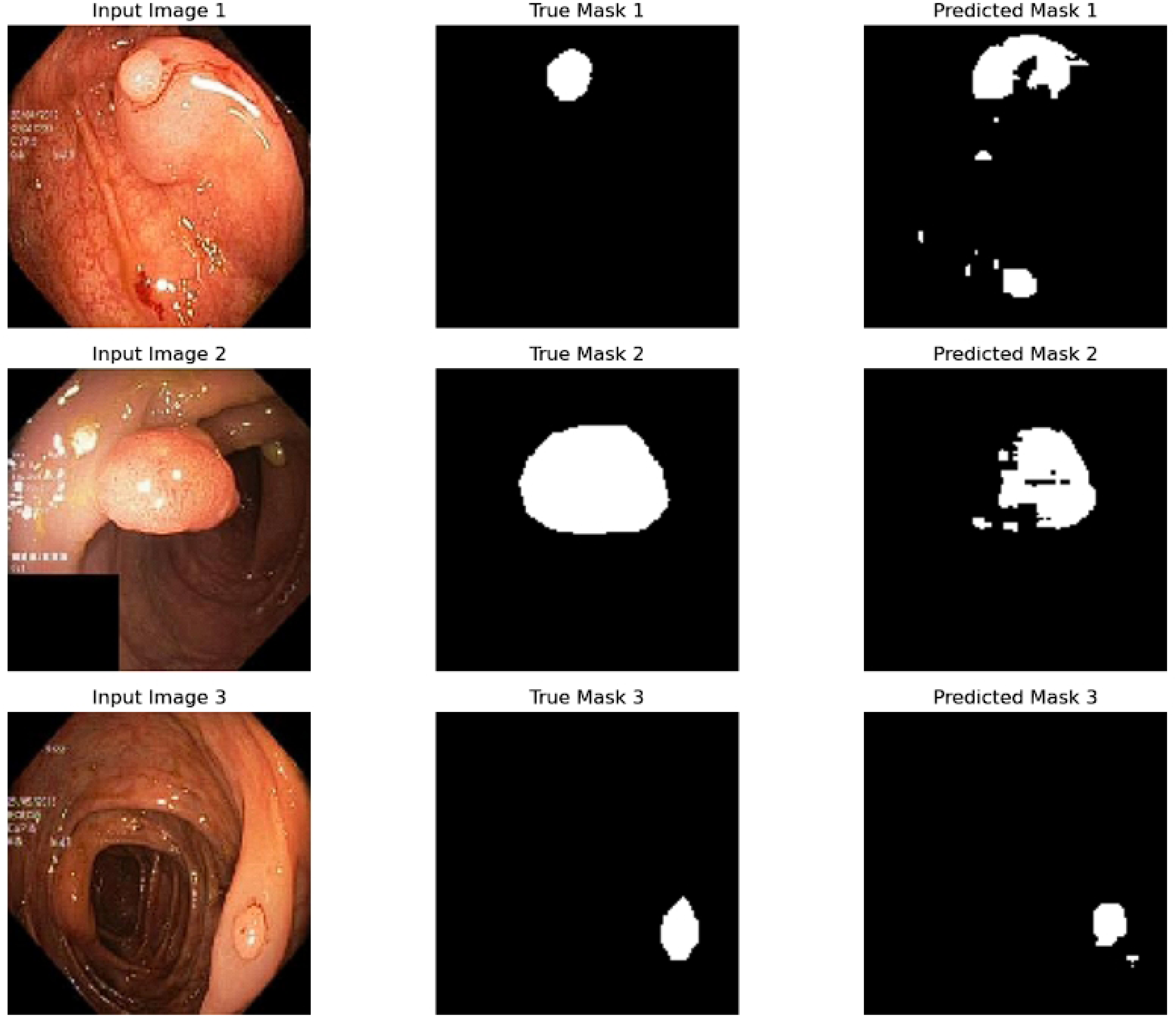
This visible image showcase the input images with their true masks and predicted masks, revealing a notable leap from decent to superior segmentation precision as the training dataset scales up from 1000 to 1800 images for the SegNet model.

The result of the SegNet model shows a comparison between true masks and predicted masks for three input images. In the result, the true masks accurately outline the regions of interest, while the predicted masks show varying degrees of accuracy. For example, the first input image of the prediction after training with 1000 images in (a) of Figure 9,the predicted mask captures the shape perfectly. For input image 2, the predicted mask is relatively close to the true mask, but misses some details. And last of all, for the input image 3, the predicted mask aligns well with the true mask in shape but has minor inaccuracies which includes some extraneous areas. Overall, the model performs reasonably well with 1000 images (without any negative images), although refinement could improve precision. Again, in this section, the true masks accurately delineate polyp regions as white areas with a black background. The predicted masks for the first input image in (b) of Figure 9, show the predicted mask captures the general shape but includes extra areas, and irregular boundaries compared to the true masks. For the second input image of (b) in Figure 9 the predicted mask resembles the true mask, but there are some missing details. Again, for the third input image of (b), the predicted mask approximates the shape but lacks precision. The inclusion of 800 negative images seems to influence the improvement of the model’s ability to differentiate healthy tissue.

The instability of the SegNet model, combined with the consistent increase in training precision, suggests potential overfitting due to the limited size of the dataset or the complexity of polyp segmentation, as observed in studies on medical image segmentation [39]. These challenges are particularly noticeable in the SegNet sensitivity to irregular polyp shapes, which complicates convergence. Data enhancement techniques were explored to reduce this issue, but further optimization is needed to enhance model stability.

(a) Prediction after training with 1000 Image
(b) Prediction after training with 1800 Image

### ResNet Model Output Analysis

#### Training Behavior

Figure 10 shows the training and validation curves of ResNet on both datasets.

**Fig 10.**
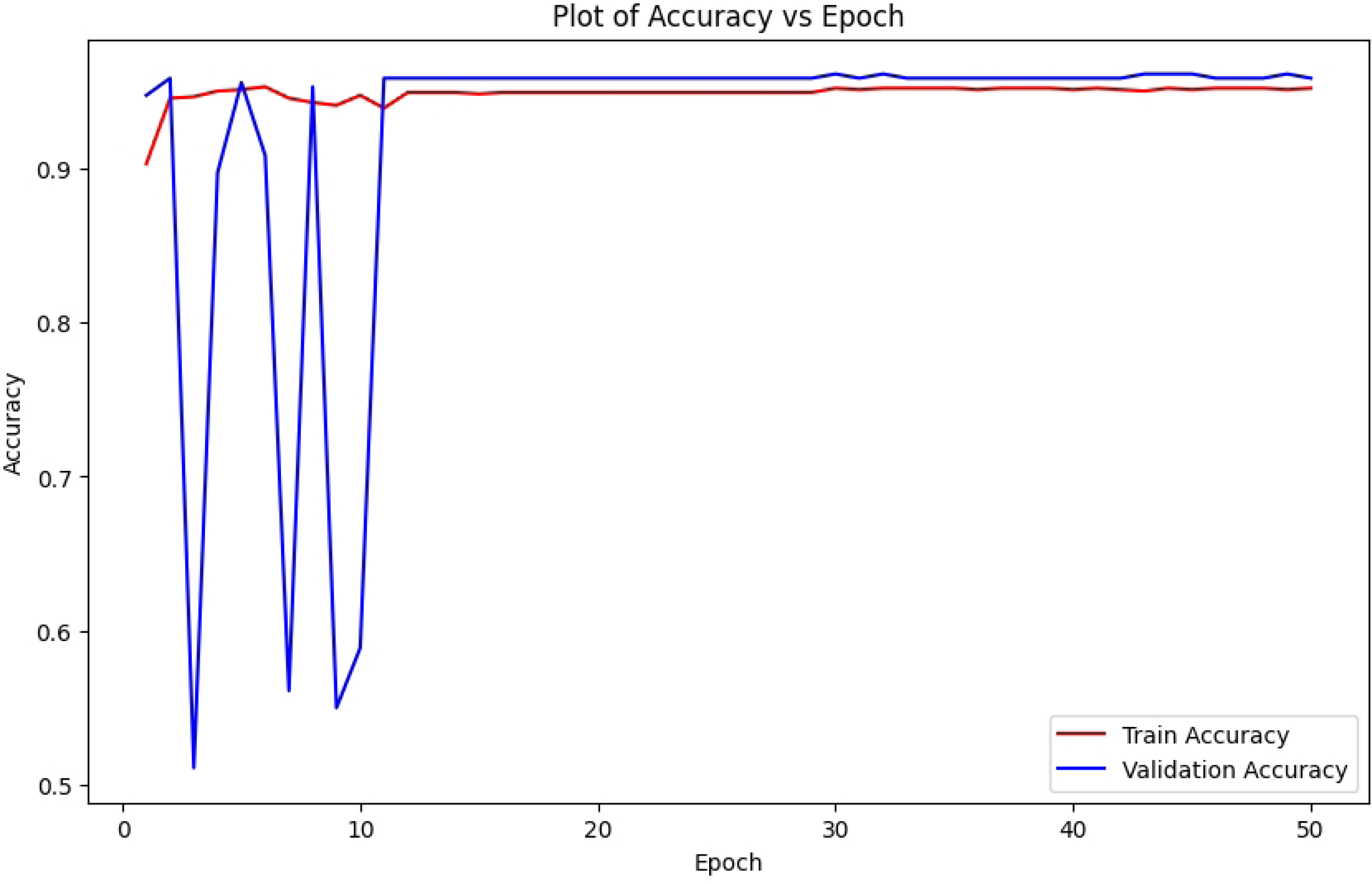
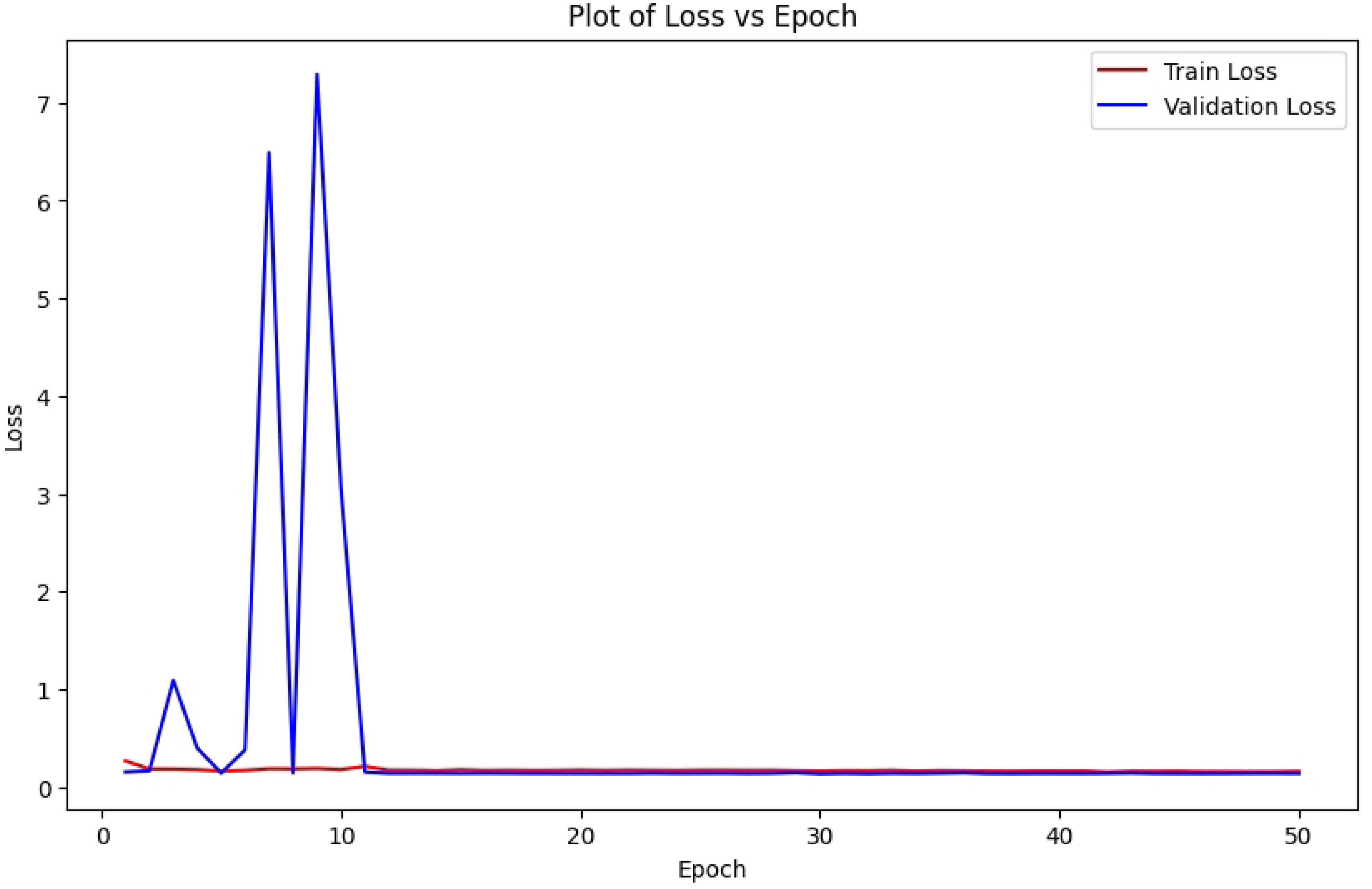
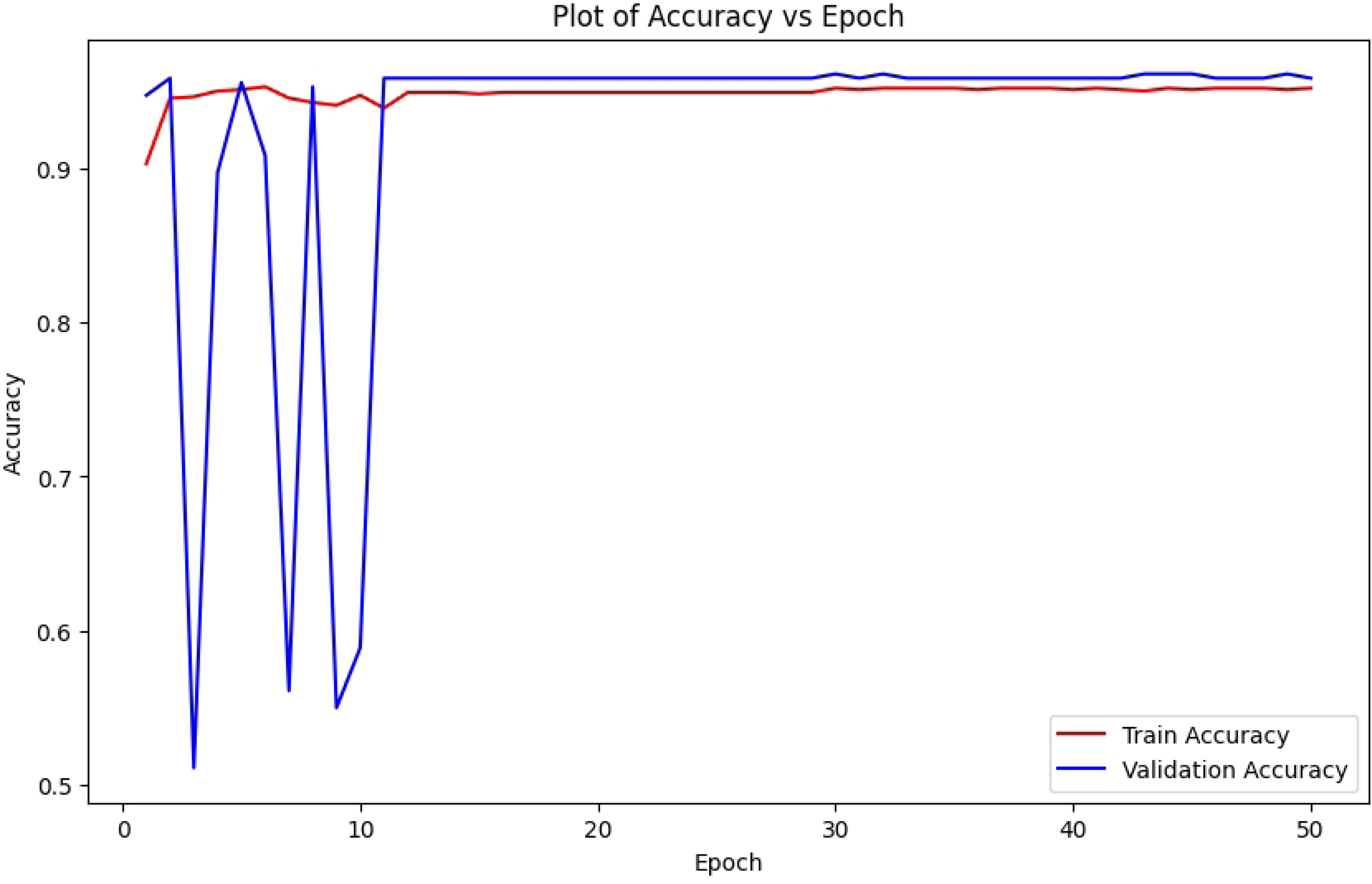
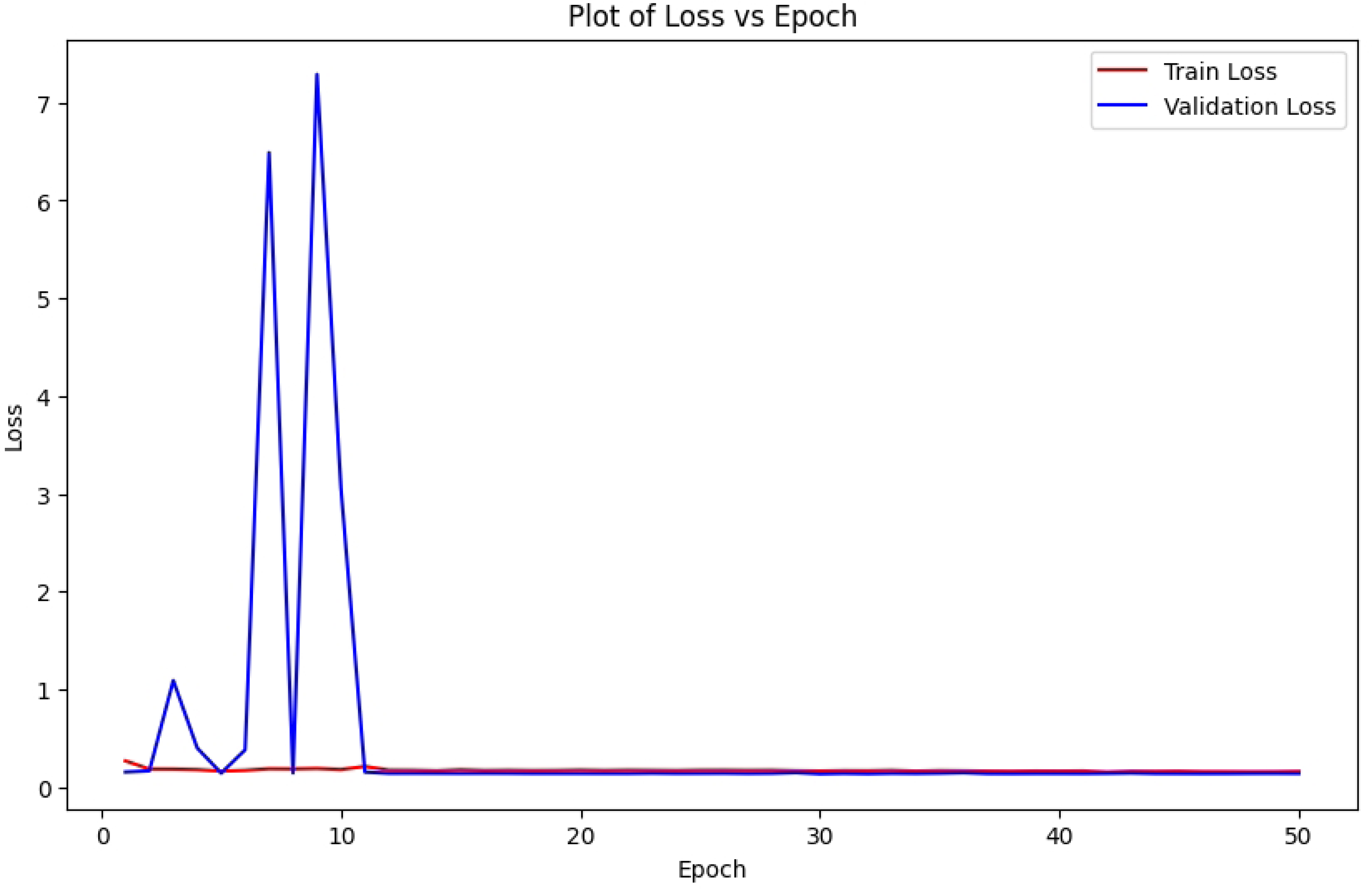
These graphs depict a swift initial increase and subsequent leveling off of accuracy, accompanied by notable early loss peaks that later stabilize, underscoring the ResNet model’s performance and possible data irregularities over 50 epochs with datasets of 1000 and 1800 images, both achieving a stable accuracy.

The two graphs represent the training progress of a ResNet model over 50 epochs, one is train and validation accuracy vs epoch (1000 Image) (a) and the other is the train and validation loss vs epoch (1000 Image) (b) in Figure 10, which illustrates the trends in accuracy and loss. In the accuracy plot, the train accuracy begins near 0.9 and remains stable, while the validation accuracy fluctuates significantly early on, dropping to around 0.5 before stabilizing close to 0.9 after 10-15 epochs, indicating improved generalization over time. In (b) of Figure 10, the train loss starts at approximately 0.5 and decreases steadily to near zero, while the validation loss starts high at 6-7 with notable peaks before declining and stabilizing near zero after 10-15 epochs, suggesting initial instability that the model overcomes. Again, the ResNet model provided two graphs after trained with 1800 images. Among those images, 800 are negative images. The training progress of a ResNet model over 50 epochs, showcasing the trends in accuracy and loss in (c) and (d) of Figure 10. In the accuracy plot, the train accuracy starts near 0.9 and remains stable, while the validation accuracy fluctuates early on, dropping to around 0.5 before stabilizing close to 0.9 after 10-15 epochs, indicating the model improves its generalization over time. In the loss plot (b), the train loss begins at about 0.5 and decreases steadily to near zero, while the validation loss starts high at 6-7 with significant early peaks before dropping and stabilizing near zero after 10-15 epochs, suggesting the initial instability that the model overcomes. Overall, the ResNet model demonstrates effective learning, with converging train and validation metrics after an initial adjustment period, indicating a well-trained model with good generalization and minimal overfitting.

(a) Train and Validation Accuracy VS Epoch (1000 Image)
(b) Train and Validation Loss VS Epoch (1000 Image)
(c) Train and Validation Accuracy VS Epoch (1800 Image)
(d) Train and Validation Loss VS Epoch (1800 Image)

### Qualitative Output

Figure 11 presents qualitative outputs from ResNet before and after adding negative data. Each sample includes:

**Fig 11.**
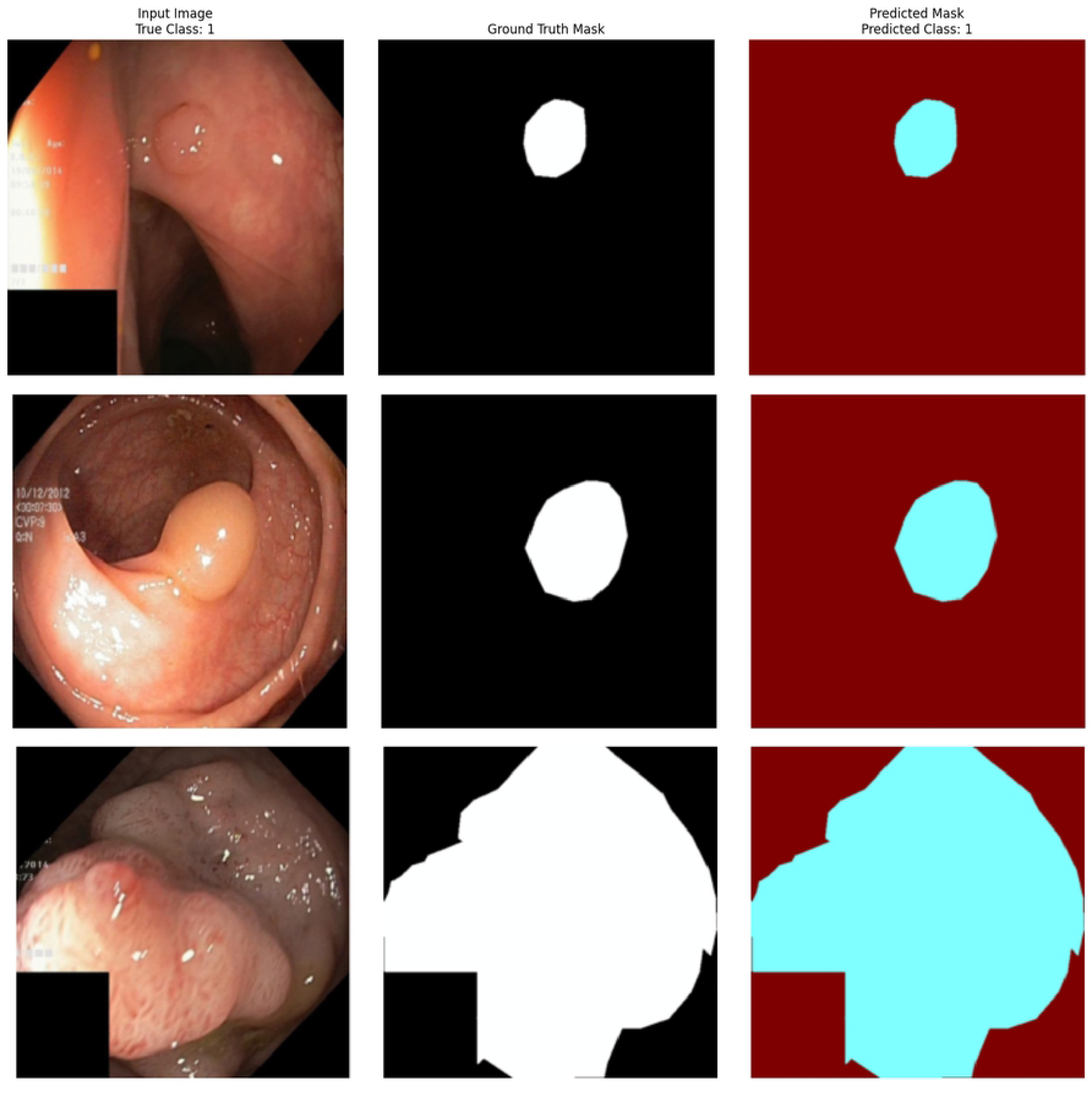
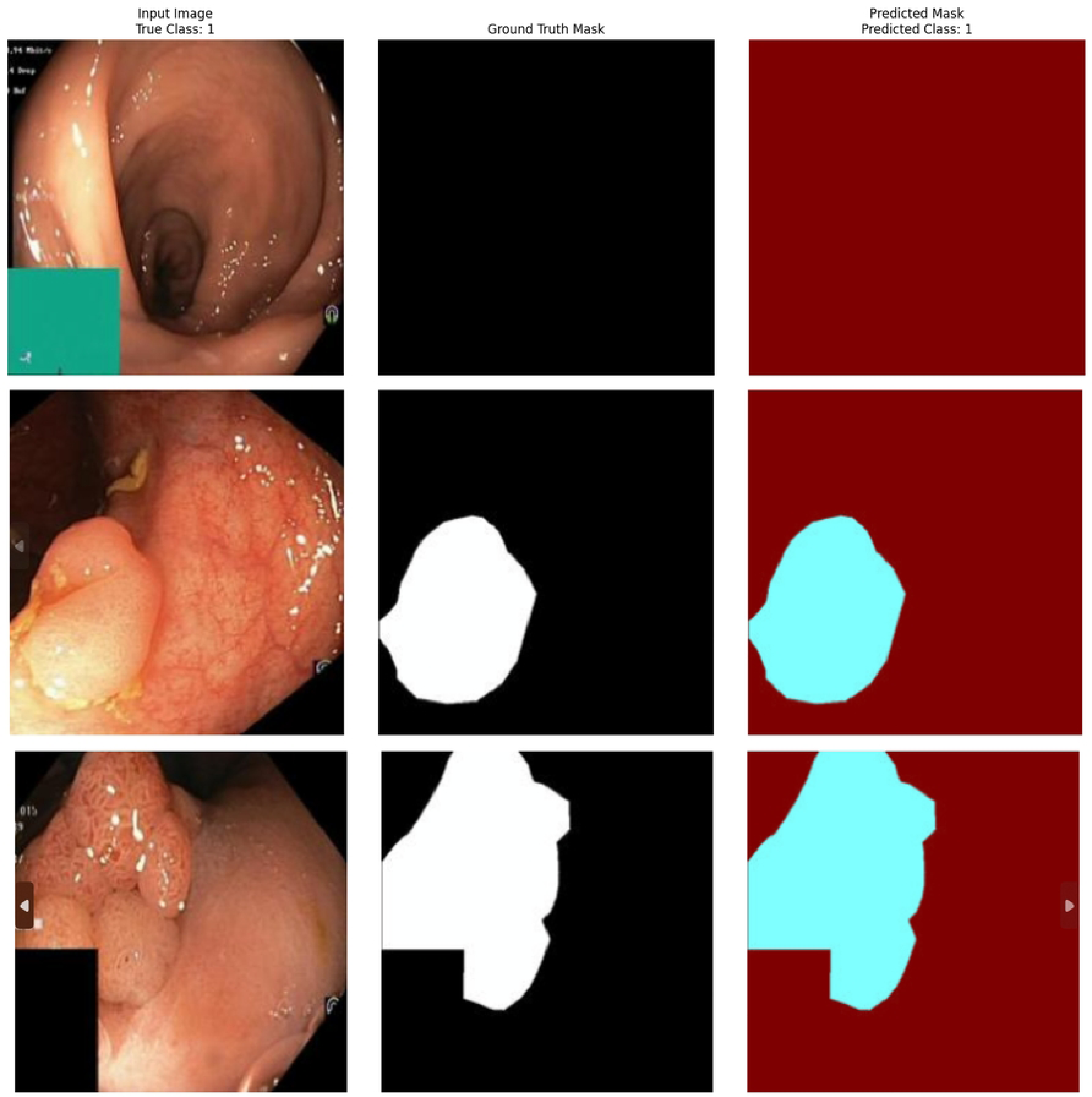
This figure presents the input images alongside their corresponding ground truth and predicted masks, highlighting a clear improvement in segmentation accuracy as the training dataset grows from 1000 to 1800 images for the ResNet model. The result highlights how increasing dataset size enhances the model’s capability to accurately identify the polyp shapes and boundaries.

The predicted masks in Figure 11, illustrate the reasonable alignment with the ground truth masks, effectively capturing the general shape and location of the polyps. For the first image of (a), which is trained with 1000 images without any negative images, the prediction closely matches the ground truth. But it has slight boundary noise. In the second image of (a) in Figure 11, shows a well-defined polyp prediction, but the shape is less precise compared to the ground truth. The third image reveals a larger inconsistency, with the predicted mask overestimating the polyp size and including extraneous regions. For (b) in Figure 11, the predicted masks show a reasonable estimation of the ground truth masks for polyp regions, with the first two images exhibiting good shape similarity, though with some boundary noise and slight over-segmentation. The third image also does not show any polyp, the ground truth mask and predicted mask demonstrate a closer match, likely because of the learning from the inclusion of 800 negative images that enhance the model’s ability to differentiate healthy tissue.

The ResNet model indicates effective learning, with converging training and validation metrics after an initial adjustment period, which also demonstrates good generalization and minimal overfitting, consistent with its robustness in medical imaging related tasks [40].

(a) Prediction after training with 1000 Image
(b) Prediction after training with 1800 Image

### Summary of Results

The addition of negative data contributed to more stable training, leading to cleaner and more accurate precise prediction outputs. Quantitative results showed moderate to substantial improvements, particularly in decreasing false positives. While U-Net performed with high accuracy, but struggled in handling polyps with irregular shapes. Meanwhile, SegNet displayed overfitting tendencies, restricting its ability to generalize effectively. In contrast, ResNet stood out for its stable generalization and minimal overfitting, which makes it well suited to a broad range of clinical scenarios. Overall, all these results highlight the necessity of selecting models according to both dataset complexity and clinical requirements.

## Discussion

This study examined deep learning models like U-Net, SegNet and ResNet for the task of colorectal cancer image segmentation. It is with particular attention to how the inclusion of negative samples influences model performance. Among the three, U-Net consistently delivered the strongest results as it was achieving a validation accuracy of 95% and demonstrating stable learning behavior. Its encoder–decoder framework strengthened by skip connections which allowed the model to retain fine spatial details as a result accurately define the boundaries of colorectal abnormalities. This aligns with the findings of Ronneberger et al. [10]. This performance is on par with Jha et al. [17], earlier findings that highlight U-Net’s ability to achieve high segmentation accuracy in biomedical application even with relatively limited training data. Moreover, SegNet showed inconsistent validation outcomes with fluctuating accuracy and loss curves. This instability indicates that the model struggles to generalize particularly when faced with more complex segmentation tasks. This result is consistent with Badrinarayanan et al. [6]. These limitations are linked to its dependence on max-pooling indices which often reduces spatial precision. ResNet displayed strong feature extraction capabilities and was more stable in terms of generalization. However, it showed signs of overfitting and was less effective for fine-grained, pixel-level segmentation tasks. This aligns with observations by He et al. [15]. These all echo earlier concerns that deep ResNet structures require careful regularization when applied to small medical datasets. A major outcome of this work was the positive impact of including negative samples. This step enhanced the model’s ability to differentiate healthy from abnormal tissues which significantly reduced false positives and improved overall specificity, a finding supported by Zhang et al. [29]. All this reinforces previous research stressing the importance of balanced datasets for medical image analysis. To summarize the whole study we can say the results highlight U-Net’s particular suitability for handling the complex textures and structural variations typical in colorectal images. Furthermore, recent advances in attention-based mechanisms, as explored by Chen et al. [30], transfer learning, as investigated by Shin et al. [31] and lightweight architecture, as proposed by Li et al. [32] or hybrid model designs, as proposed by Wang et al. [33] suggest promising directions to further strengthen segmentation performance. In clinical practice, these improvements could enhance diagnostic reliability, streamline workflows as a result ultimately support better patient outcomes.

### Conclusion

This study presented a comparative evaluation of deep learning models for colorectal cancer image segmentation using datasets that included both positive and negative samples. Therefore, aligning the evaluation more closely with real-world clinical conditions. Among the models, U-Net emerged as the most accurate. However, it encountered some challenges when dealing with highly irregular polyp shapes. Moreover, SegNet demonstrated instability and signs of overfitting. Lastly, ResNet generalized better but lacked U-Net’s precision in boundary detection. The findings emphasize the need to align model selection with clinical requirements because each architecture offers distinct strengths and limitations. Future research should focus on combining complementary models into hybrid frameworks alongside exploring more advanced data augmentation strategies. These will leverage domain-specific pretraining to enhance performance. Thus, integrating multimodal imaging data and optimizing models for real-time use may further increase their clinical applicability. This has led to the development of more effective and reliable diagnostic tools for detecting colorectal cancer.

## Acknowledgments

The authors would like to express their sincere gratitude to the Biomedical Sciences and Engineering Reseach Center (BIOSE), BRAC University, for providing the necessary resources, guidance, and support throughout the course of this research. The facilities and academic environment at BRAC University have played a crucial role in enabling the successful completion of this study.

